# Mouse α-synuclein fibrils are structurally and functionally distinct from human fibrils associated with Lewy body diseases

**DOI:** 10.1101/2024.05.09.593334

**Authors:** Arpine Sokratian, Ye Zhou, Meltem Tatli, Kevin J. Burbidge, Enquan Xu, Elizabeth Viverette, Addison M. Duda, Yuan Yuan, Samuel Strader, Nirali Patel, Lauren Shiell, Tuyana Malankhanova, Olivia Chen, Joseph R. Mazzulli, Lalith Perera, Henning Stahlberg, Mario Borgnia, Alberto Bartesaghi, Hilal A. Lashuel, Andrew B. West

**Author notes:** **Author Contributions:** A.S., Y.Z., M.T., K.B., E.X., E.V., A.M.D., Y.Y., S.S., N.P., L.S., T.M., O.C., and L.P. performed research; A.S., Y.Z., M.T., K.B., E.X., E.V., A.M.D., J.M., L.P., H.S., M.B., A.B., H.L, and A.W. designed research; A.S., Y.Z., M.T., K.B., E.X., A.M.D., N.P., J.M., L.P., H.S., M.B., A.B., H.L, and A.W. analyzed data and wrote the paper.

## Abstract

The intricate process of α-synuclein aggregation and fibrillization hold pivotal roles in Parkinson’s disease (PD) and multiple system atrophy (MSA). While mouse α-synuclein can fibrillize *in vitro*, whether these fibrils commonly used in research to induce this process or form can reproduce structures in the human brain remains unknown. Here we report the first atomic structure of mouse α-synuclein fibrils, which was solved in parallel by two independent teams. The structure shows striking similarity to MSA-amplified and PD-associated E46K fibrils. However, mouse α-synuclein fibrils display altered packing arrangements, reduced hydrophobicity, heightened fragmentation sensitivity, and evoke only weak immunological responses. Furthermore, mouse α-synuclein fibrils exhibit exacerbated pathological spread in neurons and humanized α-synuclein mice. These findings provide new insights into the structural underpinnings of α-synuclein pathogenicity and emphasize a need to reassess the role of mouse α-synuclein fibrils in the development of related diagnostic probes and therapeutic interventions.

## Introduction

The accumulation of aggregated and fibrillar forms of α-synuclein (α-syn) in neurons is a defining hallmark of several neurodegenerative diseases that include Parkinson’s disease (PD), multiple system atrophy (MSA), and dementia with Lewy bodies (DLB), which are collectively referred to as synucleinopathies (*1*, *2*). Recent advances in cryo-EM have enabled unprecedented insight into the structural features of α-syn fibrils from different synucleinopathies, revealing disease-specific fibril structural features (*3–6*). Furthermore, *in vitro* recombinant α-syn can self-assemble into fibrils of distinct structural features, depending on the buffer composition, pH and incubation parameters (*7*, *8*). Altogether, these observations demonstrate that α-syn is capable of forming different types of fibrils. However, our understanding of the molecular and cellular determinants of fibril formation and structure in the brain remains incomplete. Increasing evidence from studies investigating the pathogenic properties of recombinant and brain-derived α-syn fibrils suggests that the molecular architecture of folds and stacking arrangements within the fibrils, as well as their biochemical properties (e.g. post-translational modifications) are key determinants of their pathogenicity and spreading patterns (*6*, *9*, *10*). The fibril strain-hypothesis postulates these differences may contribute to the neuropathological and clinical heterogeneity of PD and other synucleinopathies (*3*, *4*, *11*, *12*).

The formation and propagation of α-syn pathology occurs through recruitment of non-fibrillar (e.g., monomeric) α-syn protein into preformed or new-born fibrils in a self-propagation mechanism (13–15). α-Syn fibril formation is driven by extensive hydrogen bonding interactions involving a large segment of the protein that is highly structured and forms the core of α-syn fibrils (*3*, *16*, *17*). Different α-syn fibrils have been described with remarkable variations of β-strand folding, rotational symmetry, and helical twists, as characterized by cryo-EM structure analysis. Fibrils have been purified for structural analysis directly from postmortem brain tissues in the presence of detergents, or generated detergent-free through recombinant α-syn protein incubated with patient biofluids and tissues in seeding-aggregation assays, or produced using recombinant proteins under spontaneous aggregation reactions (*3*, *4*, *18–20*). The majority of known α-syn fibrils produced under physiological-like conditions consist of two protofilaments connected by salt bridges and intermolecular hydrophobic interactions.

Recent analyses of disease-associated *ex vivo* α-syn fibrils extracted from MSA brains (i.e., MSA-fold fibrils) revealed fibrils with core structures that are distinct from those found in PD, PD-Dementia (PDD), and DLB (*3*, *4*). Sarkosyl-insoluble α-syn fibrils extracted from MSA brain homogenates contain two types of distinct structures, each consisting of two asymmetric protofilaments each with 9 to 12 β-strands (*3*, *21*). In contrast, the proposed Lewy-fold associated with PD and DLB is composed of a single protofilament containing nine β-strands, with a presumed salt bridge (E35-K80) involved in the compact packaging of the singular subunit (*4*). However, attempts to replicate the structure of brain-derived α-syn fibrils *in vitro* through manipulating the aggregation conditions of recombinant α-syn proteins, or the use of brain-derived fibrils as seeds, have not yet been successful (*22–24*). For example, the structure of recombinant α-syn fibrils amplified from seeding with MSA-fibrils were markedly different from all synucleinopathies *ex vivo* fibril variants. Interestingly, fibrils seeded from MSA fold type II, referred to as MSA-amplified, resemble recombinant α-syn fibrils with the PD and DLB-associated E46K mutation in *SNCA* (*23*, *25*). Despite the structural heterogeneity, many types of recombinant α-syn fibrils have been consistently shown to exhibit high seeding efficiency when added to cells and primary neuronal cultures or injected into the brain of rodent and non-human primate models of α-syn pathology formation and spreading (*13–15*, *26–28*). These properties of preformed fibrils (PFFs) enabled for the first time the induction of the aggregation, fibrillization, and formation of Lewy-body like inclusions in the absence of α-syn overexpression (*26*, *28–34*). This has led to the emergence of α-syn PFF seeding based models as the most commonly used preclinical models to investigate the mechanisms of α-syn pathology formation and spreading and to validate novel therapeutic targets or anti-α-syn aggregation therapeutic strategies.

Different types of α-syn fibrils have been used as PFF seeds based on wild-type human and mouse recombinant α-syn as the major PFFs used by most laboratories. However, human α-syn fibrils, whether recombinantly expressed or purified from human tissues, poorly seed pathology in rodents and fail to produce progressive dopaminergic neurodegeneration when injected into wild-type mice and rats (*27*, *35–38*). In contrast, mouse α-syn fibrils exhibit higher seeding activity and induce more pathological spreading in rodents (*27*, *36*, *38*). For this reason, the injection of recombinant mouse α-syn fibrils are commonly used in both mice and rat studies. Mouse PFFs induce a sporadic Lewy body disease phenotype, including progressive dopaminergic neurodegeneration in combination with α-syn inclusion formations (*13*, *39–46*). Interestingly, mouse α-syn fibrils bind thioflavin T less compared to human fibrils and exhibit distinct morphological features suggesting the presence of different structural conformations (*35*, *47*, *48*). The primary sequence of mouse α-syn differs from the human ortholog in seven amino acid positions (*49*, *50*). Several reports have demonstrated that a single substitution of mouse α-syn in the non-amyloid-component (NAC) or pre-NAC domain (i.e., T53A or N87S) is sufficient for mouse α-syn to template human α-syn monomeric protein *in vitro* to the same extent as wild type human α-syn (*35*, *47*, *51*). Solid-state NMR studies identified probable structural differences between mouse and human α-syn fibrils but did not identify the nature of these differences (*52*). In complement, introduction of the S87N mutation into human α-syn increases the levels of α-syn seed-induced pathology in primary mouse hippocampal neurons (*35*). However, the sequence, molecular, and structural factors underlying differences in morphological and seeding properties of human and mouse α-syn fibrils have remained poorly understood.

Therefore, it has remained unclear whether the structure of mouse fibrils commonly used in preclinical models of PD and synucleinopathies matches to known recombinant or brain-derived human α-syn fibril structures. This knowledge gap has significant implications for the field as the mouse PFF seeding models are increasingly used to decipher the role of α-syn in the pathogenesis of PD to test and validate novel targets and therapies, often with the assumption that the mouse fibrils form structures that resemble those formed by human α-syn. To address this knowledge gap, we sought to determine the structure of mouse and human α-syn fibrils using cryo-EM under identical conditions. Our results show that mouse α-syn fibrils are composed of a single homogenous and reproducible fibril type that bears a strong similarity to human α-syn recombinant fibrils templated from MSA fold type II tissues and E46K-mutated fibrils. Furthermore, the structure of the mouse α-syn fibrils do not resemble any of the structures of disease-associated brain-derived α-syn fibrils. Interestingly, the functional properties associated with mouse α-syn fibrils are also strikingly different from human α-syn fibrils in nearly all pathological endpoints that were measured. Our findings provide novel structural insights that explain the high seeding efficiency of mouse α-syn PFFs in mouse neurons and rodents and underscore the critical importance of developing disease-relevant human α-syn expressing models of pathology formation and spreading. Models with better structural and functional overlap with disease seem achievable and essential to decipher disease-relevant pathogenic processes that increase the probability of success in translating findings from preclinical models to the clinic.

## Results

### Mouse α-syn form a core structure distinct from both recombinant and brain-derived human α-syn fibrils

To investigate and compare the structural properties of mouse and human α-syn fibrils, we purified both recombinant mouse and wild-type human α-syn proteins (Fig. 1a) from *E. coli* and induced their aggregation under conditions (endotoxin-free phosphate buffered saline, pH 7.4) that have been optimized and standardized for the production of mouse α-syn fibril preparations across different laboratories (*15*, *44*). In both mouse and human α-syn fibril preparations, cryo-EM and TEM micrographs revealed homogenous unbranched structures with twisted morphology (Fig. 1b). 2D class averages reveal a single dominant structural class for both mouse and human α-syn fibrils (Table 1, Table S1). Helical reconstructions resolved a 3.1-Å map for mouse α-syn and a 2.7-Å map for human α-syn fibrils produced under the same conditions. Cross-section projections of both fibrils showed two-protofilament architectures (Fig. 1c). Interestingly, human α-syn formed fibrils with a core structure that is very similar to that of the previously reported 6sst human wild-type α-syn conformation (*18*), whereas the high-resolution fitted atomic model for mouse α-syn revealed a very different fibril core structure forming antiparallel pairs (Fig. 1d-f). Mouse α-syn fibrils display a left-handed internal pseudo C2-symmetry with protofilaments connected through a salt-bridge formed by residues K45 and E46 (Fig. 1e). The monomer subunits stack along the fibril axis with a helical rise of 4.84-Å and helical twist of 0.84° (Fig. 1f). No other fibril structures were identified in either mouse or human α-syn fibril preparations.

**Fig. 1.**
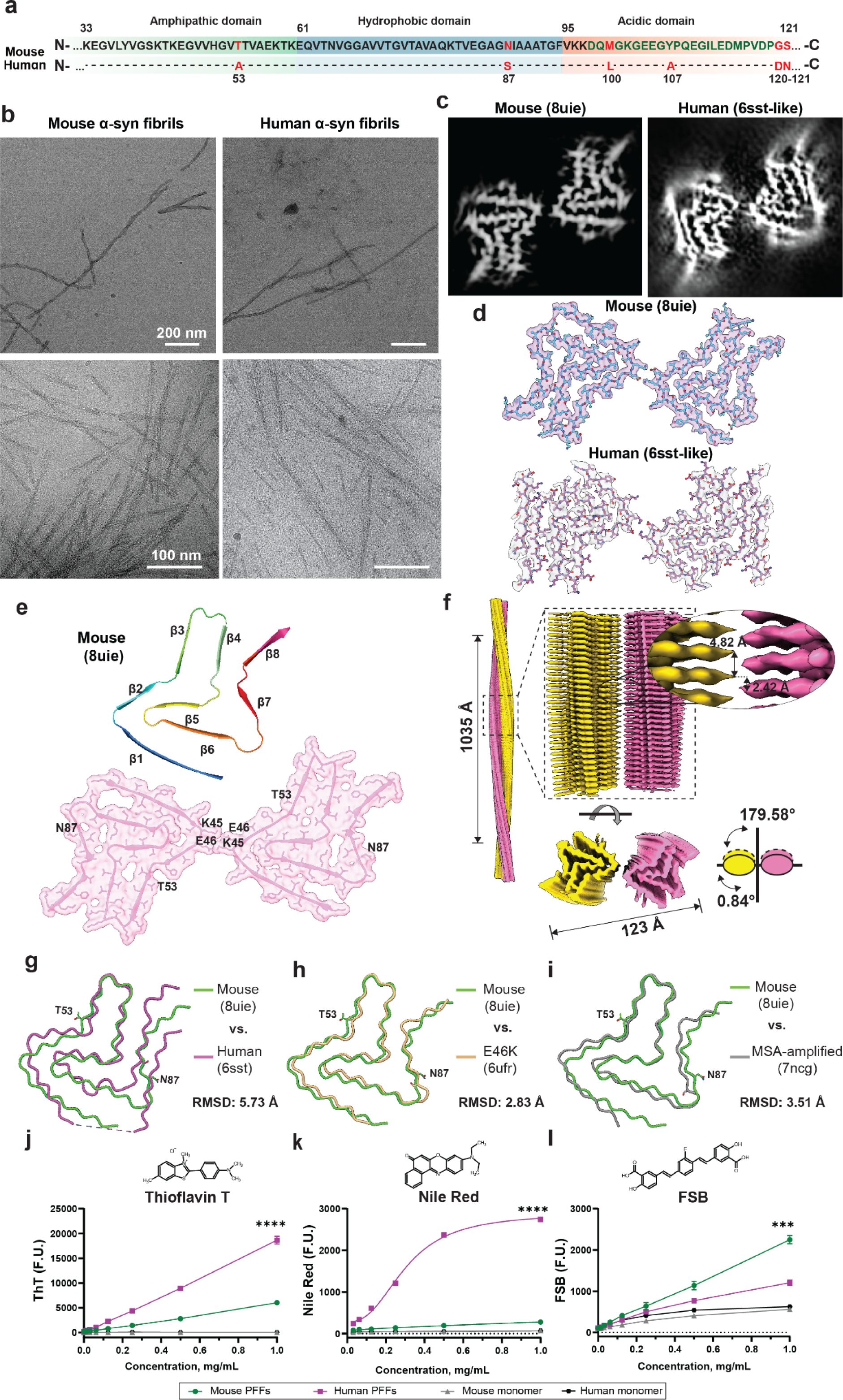
Mouse α-syn fibrils exhibit structural similarities with human E46K-mutated and MSA-amplified α-syn fibrils, yet show distinct characteristics from WT human α-syn fibrils. (a) Schematics of primary amino acid sequence of mouse and human α-syn at denoted region from 33 to 122 with indicated divergent residues highlighted in red. Depicted three domains are amphipathic in green, hydroponic in blue and acidic in orange. (b) Representative TEM (top row) and cryo-EM (bottom row) micrograph of mouse (left panel) and human α-syn fibrils with an indicated scale bars of 200 nm and 100 nm accordingly. (c) Central slice of the 3D cryo-EM map of mouse (left) and human (right) α-syn fibrils. (d) Cryo-EM density map surfaces of mouse (top) and human (bottom) α-syn fibril single filament. (e) Cartoon view of the fitted atomic model of the mouse α-syn structural arrangement, depicting β-strand positions (top panel) and cross-section view of the cryo-EM density map (shown in transparent magenta color) with an overlaid molecular model representation of the 3D density surface rendering of the fibril core. Highlighted are indicated residues involved in the formation of a salt bridge between protofilaments (K45-E46) and residues that differ from the human α-syn sequence in fibril core region (N87, T53). (f) Side view of the density map of mouse α-syn left-handed helices with a crossover distance of 1035 Å and a twist of −179.58°. The densities of the two intertwining protofilaments are colored yellow and magenta. Overlay of one layer of mouse α-syn (8uie, green) and (g) recombinant human (6sst, magenta), (h) recombinant E46K (6ufr, yellow), and (i) MSA-amplified (7ncg, gray) structures with visualized T53 and N87 amino acid schematics and indicated aligned total RMSD values. The structures were aligned based on residues 54-66 selected after an initial global fit with the lowest RMSD values. Chemical structure and binding curves of ThT (i), nile red (j), and FSB (k) to mouse and human sonicated α-syn fibrils (radii: 16.68±1.44 nm and 15.14±4.02 nm, respectively, Fig. S3) and monomeric equivalents. Data points indicate the mean from three independent experiments and presented error bars are S.E.M. ****p<0.0001 and ***p<0.01 from unpaired 2-tailed t-tests.

**Table 1.** Statistics of cryo-EM data collection and structure refinement of mouse α-syn fibril preparations.

| **Fibril type:** | **Mouse (Duke)** | **Mouse (EPFL)** |
| --- | --- | --- |
| **Data collection** |  |  |
| **Microscope** | TFS GLACIOS | FEI TITAN KRIOS |
| **Image detector** | GATAN K3 (6k x 4k) | GATAN K3 (6k x 4k) |
| **Pixel size (Å)** | 1.08 | 0.878 |
| **Defocus Range (µm)** | -2.5 to -0.8 | -2.5 to -0.8 |
| **Voltage (kV)** | 300 | 200 |
| **Exposure time (s/ movie)** | 8 | 1? |
| **Number of frames** | 55 | 30 |
| **Total dose (e<sup>-</sup>/ Å<sup>2</sup>)** | 55 | 50 |
| **Micrographs** | 3,322 | 5358 |
| **Inter-box distance (pixels)** | 27 | 16 |
| **Initially extracted fragments** | 468,092 | 271,844 |
| **Segments after 2D classification** | 42,967 | 146, 075 |
| **Reconstruction** |  |  |
| **Initial model used [PDB code]** | 6ufr | 7ncg |
| **Segments after 3D classification** | 27,524 | 50, 378 |
| **Resolution after 3D refinement (Å)** | 3.3 | 3.0 |
| **Final resolution (Å)** | 3.1 | 2.9 |
| **Estimated map sharpening B-factor (Å<sup>2</sup>)** | -56 | -52 |
| **Helical rise (Å)** | 2.42 | 2.39 |
| **Helical twist (°)** | 179.58 | 179.6 |
| **Atomic model** |  |  |
| **EMDB code** | EMD-42294 | EMD-50023 |
| **PDB code** | 8UIE | 9EWV |
| **FSC threshold** | 0.143 | 0.143 |
| **Composition** |  |  |
| - Chains | 12 | 12 |
| - Atoms(non-Hydrogen) | 5352 | 5676 |
| - Residues | 768 | 768 |
| **B-factors (Å<sup>2</sup>)** |  |  |
| -Protein | 100 | 98.25 |
| **R.M.S. deviations** |  |  |
| - Bond lengths (Å) | 0.002 | 0.009 |
| - Bond angles (°) | 0.434 | 0.918 |
| **Validation** |  |  |
| - MolProbity score | 1.44 | 1.74 |
| - Clashscore | 8.10 | 7.25 |
| - Poor rotamers (%) | 0 | 0 |
| **Ramachandran plot** |  |  |
| -Favored (%) | 98.39 | 95.16 |
| -Allowed (%) | 1.61 | 4.84 |
| -Disallowed (%) | 0.00 | 0 |

In an effort to substantiate these findings, our collaborators generated an independent set of mouse α-syn fibrils (endotoxin-free phosphate buffered saline, pH 7.4) and prepared them for cryo-EM analysis at the École Polytechnique Fédérale de Lausanne (EPFL). The conditions for generating mouse α-syn fibrils were standardized across both locations, with only minor variation in the concentration of monomeric protein (refer to Table S2 for details). Collected cryo-EM micrographs highlight unbranched homology similar to that observed at the Duke site (Fig. S1a). Cross-sections obtained at both sites were similar, further validating initial results (Fig. 1c, Fig. S1b). The reconstructed cryo-EM map of the EPFL mouse α-syn (PDB: 9ewv) fibrils revealed the same structural characteristics as those observed at the Duke site (Fig. 1e-f, Fig. S1c-e). Although the collection and refinement methods at the two sites differed, including the use of different initial models (6ufr for Duke; 7ncg for EPFL), the consistency in the structures highlights the robustness of our experimental design and the reproducibility of our findings (Fig. S2a-c, Table 1). For clarity and distinction, we will refer to the 8uie mouse cryo-EM map as ‘Duke’, acknowledging its origin at Duke University, and the 9ewv map as ‘EPFL’ henceforth.

When comparing the mouse and human fibril structures, significant shifts in the β-strands are observed near areas where the protein sequences differ, specifically at residues T53 (A53 in humans) and N87 (S87 in humans, Fig. 1a, g; Fig. S1d). Differences in the β-strand folding can be attributed to steric hindrance caused by N87, which is involved in forming an antiparallel β7 fold. In contrast, S87 might facilitate disruption of the β-sheet into a loop. We noticed that the β-strand positioning associated with mouse fibrils curated at both sites overlaps with α-syn fibril structures previously reported for E46K-mutated α-syn (6ufr), as well as wild-type human α-syn fibrils amplified from MSA brain-derived fibrils (7ncg, Fig. 1h,i, Fig. S1g,h and (*23*, *25*)). The alignment of mouse α-syn fibrils in a global fit of five stacked fibril rungs demonstrate a particularly close alignment to fibrils obtained from human α-syn monomers seeded with MSA-amplified fibrils (Fig. 1i, S1h, S3a,b). However, the twist length is different between mouse fibrils (1035-Å) and MSA-amplified fibrils (900-Å), as well as the twist angle (0.84°versus 0.95°, Fig. S3). Spontaneous with wild-type human and PD-associated α-syn fibrils or PTMs and seeded reactions generated under various buffer conditions showed lower compatibility when compared to Duke and EPFL mouse structures (Fig. S4, S5). These data show that mouse α-syn fibrils align with MSA-amplified fibrils, but not other human wild-type α-syn fibrils produced under commonly utilized experimental conditions. Neither mouse nor human fibril structures, produced under these conditions, aligned well with reported structures for sarkosyl-extracted α-syn fibrils from the brain (Fig. S6 and (*23*)).

Notable differences in twist, angle, and β-fold patterning between mouse and human α-syn fibrils suggest the possibility of differential interaction with amyloid dyes that bind in pockets affected by these patterns, for example thioflavin T, nile red, and the Congo Red-derivative FSB that are commonly utilized in biomarker seeded aggregation assays and assessments of α-syn pathology in human brains and preclinical models of synucleinopathies (*53–55*). To test this hypothesis, both mouse and human α-syn fibrils were fragmented by sonication to uniform populations (i.e., PFFs) and then concentrations of particles matched from multiple independent preparations (Fig. S7). As anticipated, the concentration-dependent curves of thioflavin T (ThT) binding revealed a much stronger affinity for human as compared to mouse PFFs (Fig. 1j). Surprisingly, nile red, revealed a large discrepancy between human and mouse α-syn PFFs (*56*, *57*). Nile red binds poorly to mouse PFFs but strongly to human variant, suggesting mouse α-syn fibrils do not have significant surface-exposed hydrophobic clusters (Fig. 1k). These observations are supported through the Congo red dye derivative FSB, with two distinct hydrophilic groups, which binds tighter to mouse than to human α-syn fibrils (Fig. 1l). To gain more insight into the structural-basis of the differential binding of the amyloid dyes to human and mouse α-syn fibrils, we carried out molecular dynamic analysis under physiological conditions. We generated 2D hydrophobicity projections of generated cryo-EM maps that support the high hydrophobicity associated with human α-syn fibrils and low hydrophobicity associated with mouse α-syn fibrils (Fig. S8a). *In silico* 3D models of electrostatic density showed a higher level of charged clusters in mouse α-syn fibrils which appear to correspond to the hydrophobicity projections (Fig. S8b). Altogether, these results demonstrate that human and mouse α-syn fibrils exhibit distinct structural, surface and dye binding properties. The striking similarities between the structure of the mouse α-syn fibrils and the MSA-amplified human α-syn fibrils (Fig. S5) suggest that the structure of the latter may be largely the result of the spontaneous aggregation properties associated with α-syn monomers. Finally, our findings support the utility of different amyloid-binding dyes, especially Nile red, that could be useful for optimizing seeding amplifications assays where differences in dye-binding has emerged as one of the key distinguishing features of samples isolated from different synucleinopathies (i.e. in the amplification of MSA vs PD CSF samples(*53*)).

### Mouse α-syn fibrils exhibit increased fragmentation and sensitivity to detergents

Protein fibrils tend to be very stable in pathogenic forms, but can also show a great range of stabilities, presumably due to relatively minor structural changes that associate with disease in complex ways (*58*). Mouse α-syn fibrils are known to be very efficient in seeding new inclusions in neurons, but also readily fragment with sonication and are poorly resistant to detergents (*59–61*). Several studies have shown that the length of α-syn fibrils is a strong determinant of α-syn seeding activity and pathology spreading in different models, with shorter fibrils (i.e., 20-100 nm) exhibiting the highest seeding activity (*46*, *62*, *63*). Fibril breakage increases the number of fibril ends, thus providing more surfaces to template the misfolding and aggregation of α-syn monomers. Furthermore, shorter fibrils are more likely to be internalized and secreted by neurons, which is expected to translate to increased seeding and spreading activities (*46*, *62*, *64*). Therefore, fibrils that are prone to fracturing may be more advantageous in spreading throughout a cell or across interconnected circuits in driving pathological properties, with fibril fragility linked to cellular cytotoxicity (*65*, *66*). The tendency of an amyloid fibril to fracture and partially denature can be influenced by the strength of β-sheet layer interactions across fibril rungs, further modified by backbone hydrogen bonding and hydrophobic interactions (*67*).

We hypothesized that the increased seeding activity of mouse fibrils could also be due to their reduced stability and high tendency to fragment. To test this hypothesis, we examined the high-resolution cryo-EM structures of both human and mouse α-syn fibrils. We noticed that mouse α-syn rungs were stacked flatly. The β-sheets deviated from a horizontal plane more than 2-Å in height only between residues 76-82 in β-strand six (Fig. 2a). In contrast, human α-syn fibril rungs are kinked and rise significantly higher along the horizontal plane in β-strands from residues 61 to 90 (Fig. 2b). Next, we conducted a computational analysis to estimate the tensile strength required to break these interactions in a predicted six-runged fibril particle (Fig. 2c). The energy produced for mouse was 1.35-fold lower than that for human α-syn fibrils in the model (Fig. 2c). Decreased energy required to disrupt the mouse fibril layers implicates a higher sensitivity to stress fracturing. To test tensile strength of fibrils during sonication, we subjected full-length mouse and human α-syn fibrils to ultrasonic waves produced by a cup-horn water bath-cooled apparatus that avoids probe-tip positioning effects to minimize heterogeneity in energies applied across the sample. Ours and others previous studies have shown that the length of terminally sonicated mouse α-syn fibrils ranges from 15-25 nm under common sonication conditions in physiological buffers, while full-length intact fibrils are much longer and exceed well over 100 nm in length (*68–70*). Approximately 31.24% (+/−2.54) of mouse α-syn fibrils, compared to 2.08% (+/−0.41) of the human conformer, fracture to less than 100 nm in size after 2 minutes of sonication, suggesting mouse α-syn fibrils are drastically less stable. When the sonication process was extended to 30 minutes, the majority of mouse fibrils (64.5%, +/−2.74) were fractured to less than 100 nm, whereas a larger proportion of longer fibrils persisted in the human fibril pool (53.77%, +/−2.92, Fig. 2d).

**Fig. 2.**
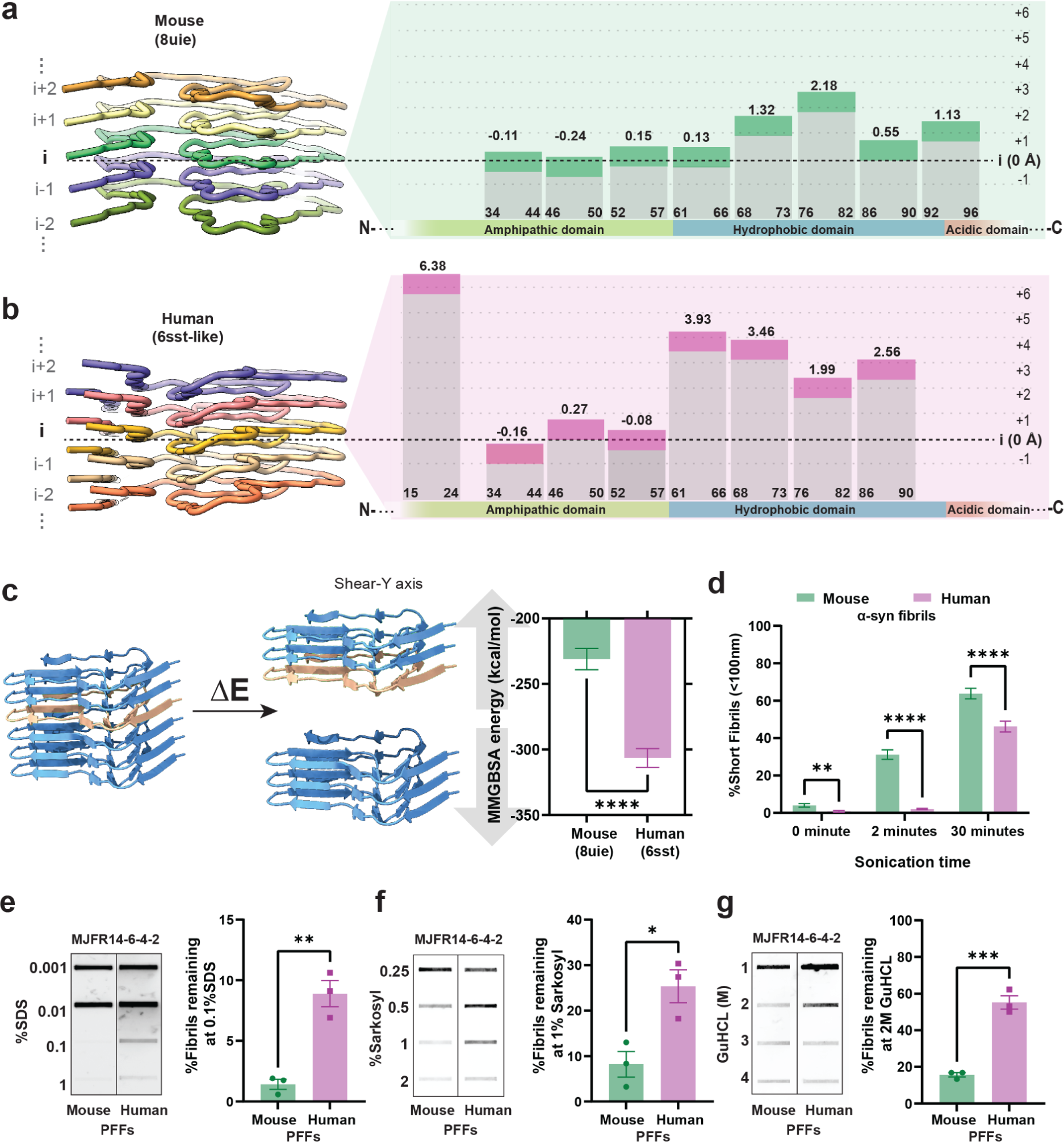
Distinct β-fold stacking arrangements in mouse α-syn fibrils contribute to low tensile strength and resilience. Cross-β-structure of five protofilament rungs (left) and schematic representation of stacking arrangement (right) with estimated coordinates against the assigned baseline position (i) as the horizontal plane (37-57 amino acids) aligned to primary peptide sequence for mouse, 8uie (a) and human, 6sst (b) cryo-EM models. (c) Representation of proposed model of β-sheet fragmentation for tensile strength estimation used to simulate the MMGBSA energy of α-syn fibrils rupture and group analysis of MMGBSA energy required to disrupt a stack of 6 protofilaments. Error bars represent SD of 100 independent simulations. (d) Group comparison of fibril breakage under sonication conditions shown as the percent of size population of 10-100 nm at 0, 2, 30 minutes measured by DLS approach. Error bars indicate S.E.M of three independent experiments with 30 acquisition measurements for each biological sample. Filter-trap slot-blot analysis of sonicated fibrils (PFFs) exposed to SDS (e), sarkosyl (f), guanidine chloride (g) concentrations detected by α-syn aggregate-specific antibody MJFR14-6-4-2 with a group analysis of the level of mouse and human fibrils detected after exposure to 0.1% SDS, 1% of sarkosyl and 2M GuHCl. Error bars in panels e, f, g indicate S.E.M from three independent experiments. *p<0.05, **p<0.01, **p<0.001, ****p < 0.0001 from unpaired 2-tailed t-tests.

Tensile strength is also linked to degradation, as fibrils that exhibit a low-force structural architecture are prone to rapid denaturation (*71*). To further assess the differential overall stability of mouse and human α-syn fibrils, we investigated their propensity to disassemble in the presence of detergents (*72*, *73*). This was achieved by monitoring the amount of remaining fibrils in the presence of increasing concentration of detergents. We conducted a filter-trap slot-blot analysis with previously fully sonicated fibrils (60 minutes, PFFs), and then incubated the PFFs with increasing concentrations of SDS, sarkosyl, and guanidine HCl to assess the amount of detergent-resistant fibrils under denaturing conditions (Fig. 2e-g). While 0.1% SDS was sufficient to denature mouse α-syn, human PFFs were significantly more resistant to denaturation, as monitored using the MJFR-14-6-4-2 monoclonal antibody, which shows higher affinity and preferential binding to aggregated forms of α-syn (Fig. 2e). Monoclonal antibody MJFR-14-6-4-2 binding intensities were normalized to fibril preparations without detergents to account for the possibility that the MJFR-14-6-4-2 antibody might have different affinity between mouse and human α-syn PFFs. Similar results, higher stability for human α-syn PFFs, were obtained using 1% sarkosyl or 2M guanidine as denaturing detergents (Fig 2f,g). Collectively, these results suggest that mouse α-syn fibrils are considerably fragile and subject to rapid fragmentation.

### Mouse α-syn fibrils have low immunogenicity

Previous studies have suggested that different human α-syn fibril structures can elicit powerful immunological responses, especially from myeloid cells like monocyte-derived macrophages (MDMs and (*74*, *75*)). Given the significant differences in the structural characteristics observed between human and mouse α-syn fibril preparations, we examined whether these differences might impact their functional interactions with immune cells. We cultured primary human MDMs from the blood of healthy individuals as previously described (*76*). Macrophages were first exposed to Alexa-657-conjugated mouse and human α-syn PFFs to determine if there were differences in cellular uptake. Both mouse and human α-syn fibrils were derived from preparations of protein processed extensively through endotoxin removal kits, and verified endotoxin free (i.e., <0.1 endotoxin units per mg of PFFs) via assessments with Limulus Amebocyte Lysate (LAL) assays. As mouse fibril preparations may present with an average shorter fibril length owing to spontaneous fracturing, immediately prior to treatment on cells, both mouse and human fibril preparations were sonicated to terminal lengths (Fig. S7). The uptake of both types of PFFs into macrophages was identical, with both fibril types equivalently localized into LAMP1-positive vesicles within minutes of addition to cell cultures (Fig. 3a-c). To investigate the inflammatory responses to the PFFs, secreted human IL-6 and CCL5, known to be robustly stimulated by human short rod-α-syn fibrils that are ostensibly similar to the 6sst fibrils used here, were measured by ELISA in a time-dependent manner similar to that previously described (*74*). With the addition of equivalent numbers of mouse and human fibril particles to the culture media, human PFFs provoked a larger IL-6 and CCL5 response than mouse PFFs (Fig. 3d,e). IL-6 and CCL5 signaling is associated with autophagy and lysosomal damage (*77*), and α-syn fibrils are thought to cause lysosomal damage in different types of cells (*78*). We noticed that internalized human PFFs tended to be co-positive in vesicles for the lysosomal damage marker Gal3, whereas few instances of internalized mouse fibril particles were associated with Gal3-positive vesicles (Fig. 3f,g). Gal3 recruitment to lysosomes has been associated with release of IL-6 upon CCL5 activation (*79*, *80*). Further analysis of colocalization with dye-quenched (DQ) mouse or human PFFs, produced as previously described to assess fibril-induced lysosomal damage, demonstrate few instances of Gal3 staining in degradative vesicles filled with mouse α-syn compared to most vesicles filled with human PFFs that show signs of damage (Figure 3f,h; (*81*)). These results suggest that human, but not mouse α-syn PFFs, damage lysosomes and cause correlated cytokine and chemokine responses. Interestingly, a previous study has shown that E46K α-syn fibrils also fail to induce lysosomal rupture compared to human isoforms (*82*). To the best of our knowledge, this is the first study that points directly to differences in α-syn fibril structure as a key determinant of α-syn disruption of lysosomes in immune cells that likely affects pro-inflammatory immunological responses.

**Fig. 3.**
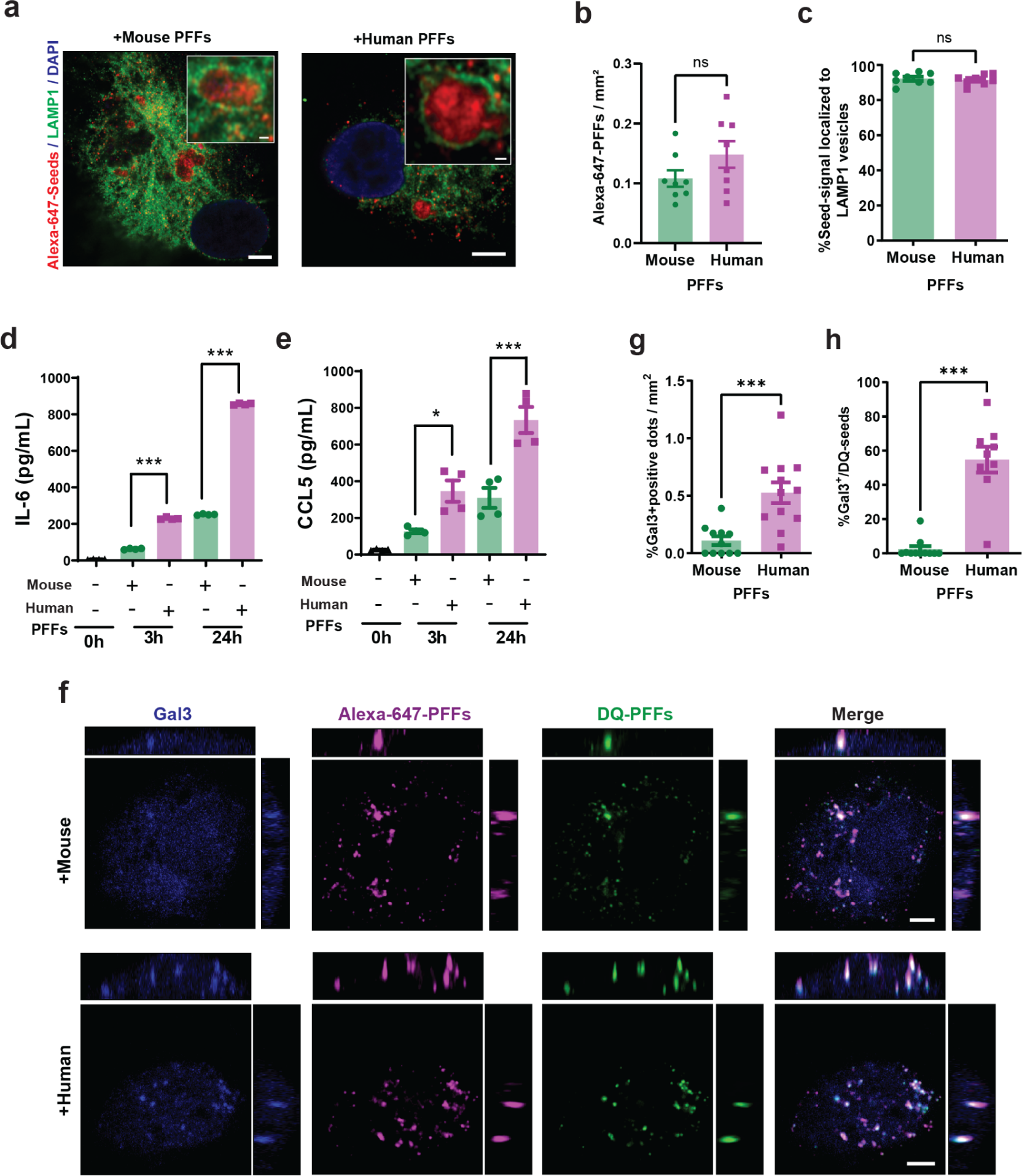
Mouse α-syn fibrils fail to elicit robust cytokine and damage responses in macrophages. (a) Representative orthogonal view of LAMP1 immunostaining after 2 hours of incubation with mouse or human α-syn PFF-treated monocyte-derived macrophages (MDMs). The magnified boxes show PFF-positive LAMP1 vesicles. The scale bar is 5 µm and 0.5 µm for the magnified boxes. (b) Number of individual Alexa-647-PFFs captured in cell area per mm². Each data point represents the mean analysis of one individual cell from three independent experiments with at least nine images analyzed per group. (c) Percentage of LAMP1-positive vesicles containing α-syn PFFs in MDM cultures treated with mouse or human PFFs. Eight images from three independent experiments were quantified for each condition, with each dot representing results from one image. ELISA quantification of the extracellular soluble IL-6 (d) or CCL5 (e) concentration in MDM cultures treated with PFFs for 3 and 24 hours. Each data point represents a signal from two technical replicates from three independent experiments. (f) Representative images of Gal3 immunostaining after 48-hour long treatment with Alexa-647- and DQ-PFFs. Orthogonal views of sequential z-stacks are shown. Side left image = *x*,*y* plane, side right image = *y*,*z* plane; top image = *x*,*z* plane. The scale bar is 5 um. (g) Percentage of Gal3-positive vesicles calculated per mm² of cell surface area in PFF-treated MDM cultures after 48 hours of incubation. Each data point represents the mean of one individual cell, from three independent experiments with at least eight images analyzed per group. (h) Percentage of DQ-PFFs in Gal3-positive vesicles in proportion to the overall DQ-fibril count after 48 hours of incubation. Each dot represents the mean analysis of one individual cell, from three independent experiments with at least eight images analyzed per group. Error bars represent S.E.M and ***p<0.001, *p<0.05 and ns for not significant from unpaired 2-tailed t-tests (panels b, c, g, h) and ANOVA with Tukey’s multiple comparisons test (panels d, e).

### Mouse α-syn fibrils are subject to clathrin-dependent endocytosis and robustly cross-seed human α-syn pathology in neurons

α-Syn fibrils are known to be robustly internalized in cells through a variety of reported endocytosis mechanisms that may vary between cell types (summarized in (*83*)). The effects of α-syn structural variation, such as that which exists between human and mouse α-syn fibrils, has not been closely evaluated on the uptake of fibrils into neurons. Differences in uptake between fibril structures could dictate downstream phenotypes such as inclusion formation and toxicity. To evaluate mouse fibril uptake, primary hippocampal neurons were cultured from human-PAC-wt-*SNCA*^+/+^/*Snca*^−/−^ transgenic mice that exclusively express the human α-syn protein at physiological levels (*68*), and low concentrations of terminally-sonicated PFFs (1 µg per mL) were added to the cultures at day-in-vitro 7. Similar to macrophages, there were no apparent differences in neuronal uptake between human and mouse fibrils labeled with pHrodo- and Alexa-Fluor 568 as monitored over a 24-hour period (Figure 4a-e). Application of the clathrin-mediated endocytosis inhibitor Dyngo 4a 30 min prior to addition of the labeled PFFs virtually eliminated any traces of mouse or human fibril uptake, whereas the 5-(N-ethyl-N-isopropyl) amiloride (EIPA) small molecule pinocytosis inhibitor had minor effects (Fig 4f,g). These results suggest that the uptake of sonicated α-syn fibrils into neurons in culture is similar between mouse and human fibrils in a clathrin-dependent endocytosis mechanism.

**Fig. 4.**
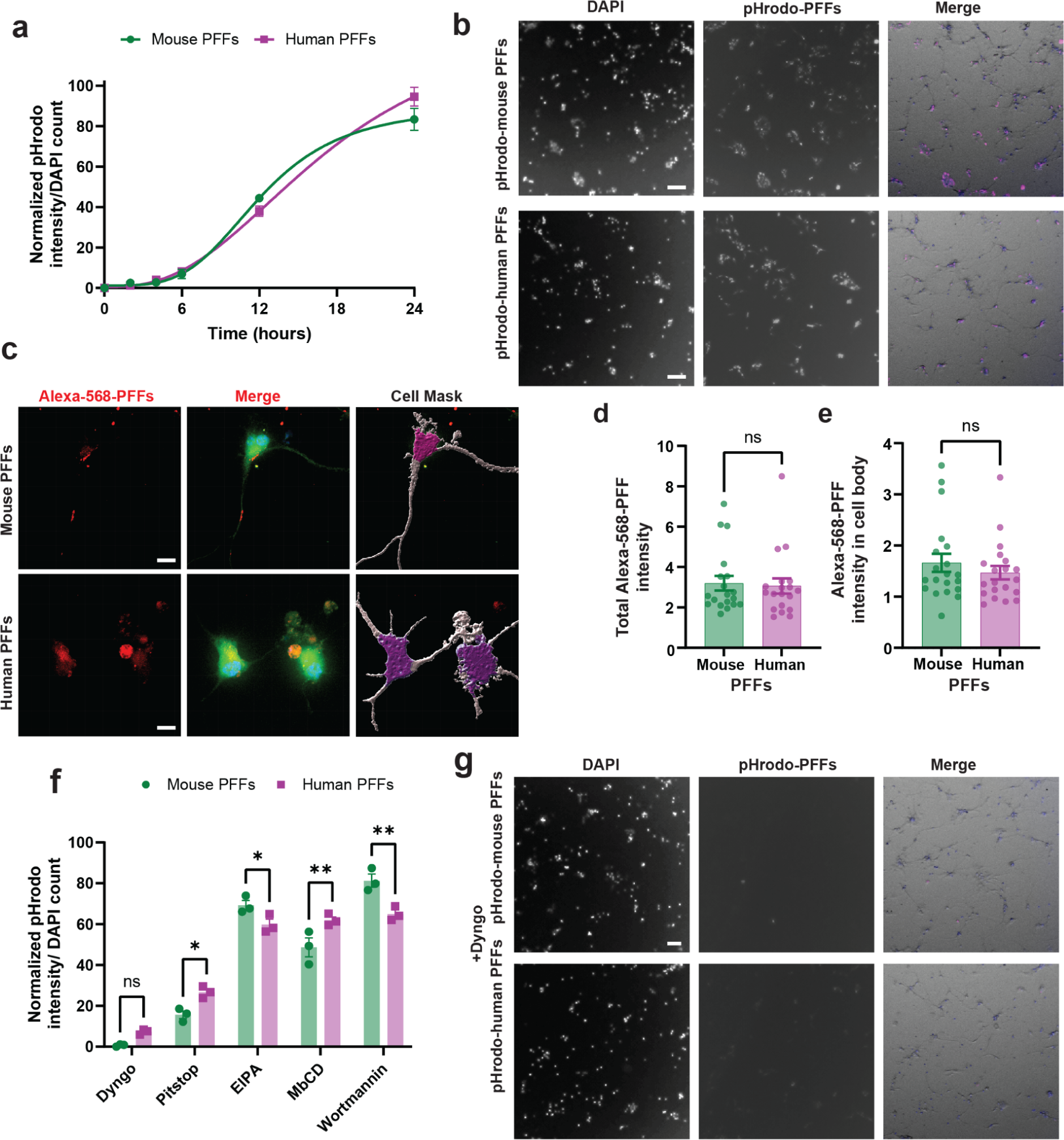
Mouse α-syn fibril uptake in neurons is clathrin-dependent and similar to human α-syn fibril uptake. (a) Time-dependent dynamics of pHrodo-labeled mouse or human PFF internalization in human-PAC-wt-*SNCA*^+/+^/*Snca*^−/−^ hippocampal primary neuron culture at DIV7 measured as normalized pHrodo-channel intensity to DAPI count. Each dot represents the mean value of three independent neuronal cultures with four images analyzed for each replicate. (b) Representative immunofluorescence images of pHrodo-labeled PFF internalization at 24 hours. Merge images incorporate overlays of phrodo-labeled PFFs signal colored in magenta, DAPI in blue, and phase-contrast in gray. (c) Representative images of ∼60-day old iPSC midbrain dopaminergic neurons treated with Alexa-568 labeled mouse or human PFFs at 10 µg/mL concentration. Merge images include Celltag labeling of total cell membrane in green, Alexa-568-labeled PFFs in red, DAPI in blue with the extracellular signal quenched using 0.1% Trypan Blue. Cell mask highlights perinuclear and neuritic areas used for the following group analysis. Group analysis of total Alexa-568 intensity in neuronal cells (d) and Alexa-568 signal in perinuclear area (e) after 8 hours of incubation with labeled α-syn PFFs. Each dot represents the mean value of one image with at least 20 images collected from three independent experiments. (f) Uptake of pHrodo-labeled α-syn PFFs at 24 hours in primary hippocampal cultures previously treated with endocytosis inhibitors shown as normalized pHrodo intensity in relation to untreated internalization rate. (g) Representative images of primary hippocampal culture at DIV7 treated with clathrin-mediated endocytosis inhibitor Dyngo for 30 min prior to pHrodo-conjugated mouse and human PFFs addition for 24 hours. Merge images include the pHrodo-PFF signal indicated in magenta, DAPI in blue and phase-contrast in gray. Each dot in each column graph represents the mean value of four images per condition from three independent experiments. Error bars for each group analysis represent S.E.M and **p<0.01, *p<0.05 and ns for not significant from 2-tailed t-tests.

With similar uptake between mouse and human fibrils, since human α-syn fibrils poorly template endogenous mouse α-syn protein (*35*), it can be reasonably predicted that mouse α-syn fibrils would also exhibit significantly reduced ability to template human α-syn protein. However, different structural and mechanical fibril properties discovered here between mouse and human fibrils that include reduced hydrophobicity and elevated rates of fragmentation may affect seeding potency in neurons. Therefore, we directly compared the seeding activity of mouse and human fibril preparations, paying close attention to equivalency in the number of fibril particles applied to cell culture models. As shown in Fig. 5, mouse PFFs were much more efficient in seeding pathology compared to human PFFs (Fig. 5a,b). Importantly, staining of neuronal cultures derived at the same time from *Snca*^−/−^ transgenic mice, which do not express α-syn, showed no evidence of pS129-α-syn pathology when treated with the same concentration of α-syn PFFs, or control monomeric α-syn (Fig. 5b, Fig. S10). Observing the pS129-α-syn patterning in neurons, there is a tendency for human PFF-seeded pathology to localize near the cell nucleus, whereas mouse α-syn fibril pathology spreads across the cell (Figure 5c-e). Other studies have likewise indicated that a higher proportion of cell body-like, or somatic, inclusion can be associated with a Lewy-body functional fibril profile, different from those found with MSA-associated fibrils that appear to spread pathology across the cell (*84*).

**Fig. 5.**
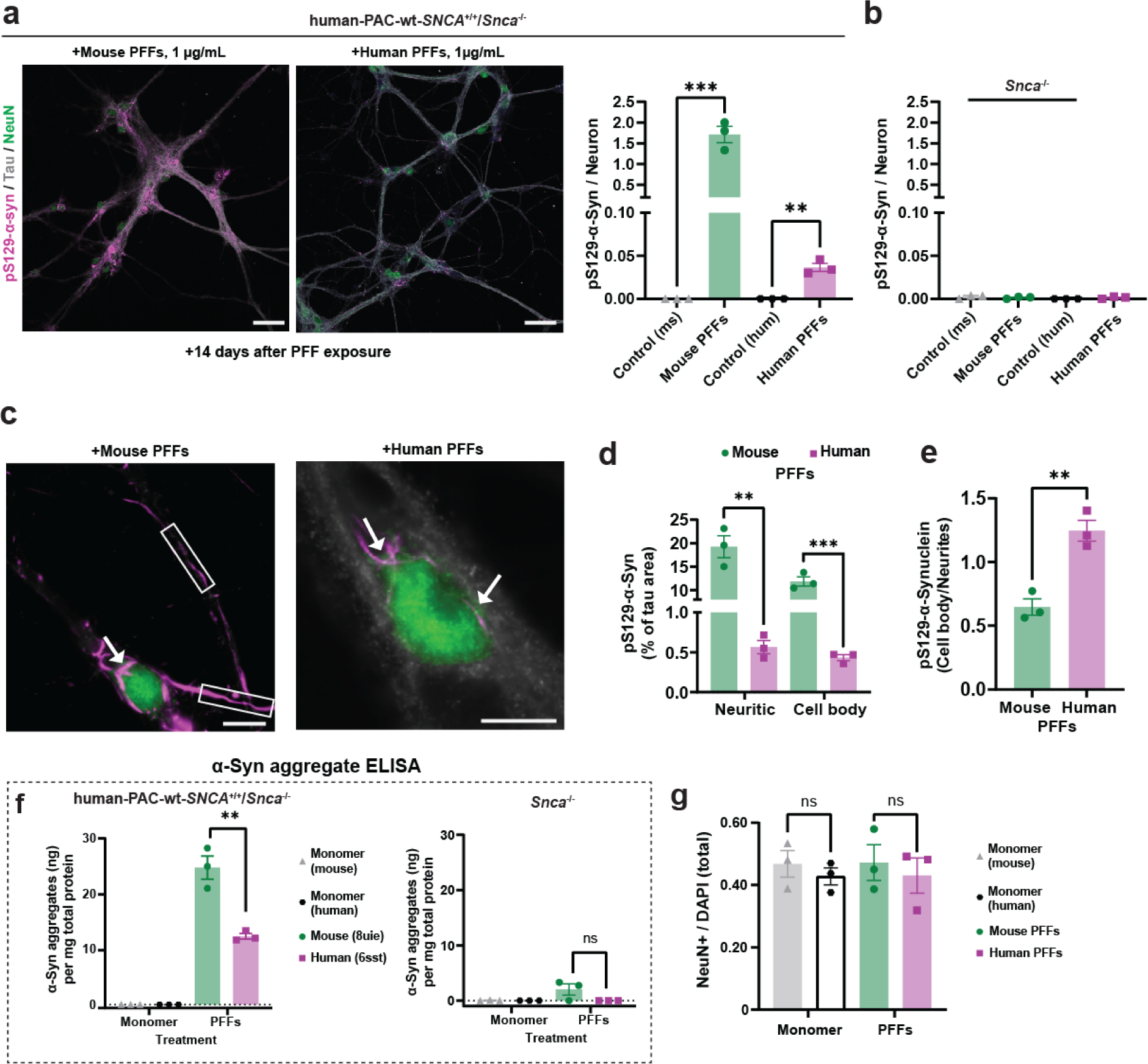
Mouse α-syn fibrils pathology propagation is more efficient than human α-syn in primary neurons. (a) Representative immunostaining of neurons cultured from human-PAC-wt-*SNCA*^+/+^/*Snca*^−/−^ hippocampal primary neuron culture treated with 1 µg/mL of α-syn PFFs for 14 days and stained against pS129-α-syn (magenta), Tau (grey) and NeuN (green) (right) and levels of pS129-α-syn signal relative to the number of neurons in corresponding cultures (left). Control groups include mouse (ms) and human (hum) α-syn monomer treatments. (b) Group analysis of pS129-α-syn signal per neuron in *Snca*^−/−^ cultures following 14 days of exposure with α-syn PFFs or monomer (ms and hum) equivalents. (c) Representative images of inclusions in the cell bodies (indicated with arrows) and neurites (outlined in square boxes) in human-PAC-wt-*SNCA*^+/+^/*Snca*^−/−^ hippocampal primary neuron culture after α-syn PFF exposure. The scale bar is 50 µm for the main image and 10 µm for magnified images. (d) Abundance of distinct pS129-α-syn signals in cell bodies or neuritic morphology in neuronal cells treated with 1 µg/mL of mouse or human α-syn PFFs. (e) Proportion of pS129-α-syn occupancy in cell body and neurites in primary hippocampal cultures incubated with mouse or human α-syn PFFs for 14 days. (f) ELISA quantification of α-syn aggregate levels in cell lysates from human-PAC-wt-*SNCA*^+/+^/*Snca*^−/−^ or *Snca*^−/−^ neuronal cultures treated with fibril PFFs or monomeric protein for 14 days. (g) Group analysis of NeuN-positive nuclei abundance normalized to DAPI count. Each data point in group analysis plots represents the mean of signal from an individual litter with two replicates per litter and at least 25 images analyzed for each replicate and error bars indicating S.E.M. Significance was determined by 2-tailed t-tests; **p<0.001, ****p < 0.0001, ns for not significant.

To further assess the extent of pathology formation, we used an ELISA designed to quantify aggregated, but not monomeric, forms of α-syn. This ELISA approach detected both mouse and human 6sst α-syn fibrils without apparent species bias down to picogram levels (Fig. S9). We found that mouse fibrils induce ∼2.5 times higher levels of α-syn aggregates compared to equivalent concentrations of human fibril particles (Fig. 5f). With this technique, we did not detect any significant signal above background in neuronal lysates from samples incubated with the same amount of monomeric α-syn protein. Further, in cultures from *Snca*^−/−^ mice (that lack any endogenous α-syn expression), incubation with either mouse or human α-syn fibrils did not yield any detectable signal after 14 days of incubation, demonstrating the ELISA does not measure the fibril particles added to the cultures after incubation, ostensibly due to their degradation in the culture over time in the absence of endogenous α-syn (Fig. 5f).

Observing the unexpected phenomenon that mouse α-syn PFFs can template significantly higher pathology in mouse neurons expressing only human α-syn, we sought to confirm these results in the context of human iPSC-derived dopaminergic neurons(iPSC-DA) prepared and cultured for 60 days by established protocol (*85*). Characterization of neuronal and DAergic markers showed that cultures contained 81% (+/− 4.1) FOXA2/TH + neurons, and 90% (+/−5%) TH/βIII-Tubulin + neurons (Fig. S11a). IPSC-neurons incubated with equivalent concentrations of Alexa-568 mouse and human PFFs for seven days revealed presence of puncta pS129-α-syn across the cell body and neurites. Notably, Alexa-568 mouse α-syn fibril particles provoked more than twice the pathology than comparable particles of human PFFs (Fig. S11b,c). Also similar to observations in mouse neurons, Alexa-568 mouse PFF-induced pS129-α-syn pathology tended to spread across the neuron through the neurite, whereas human α-syn fibril-induced pathology tended to occur predominantly in the cell body (Fig. S11d). Overall, these results demonstrate higher seeding potency of mouse PFFs and suggest that they are able to form diffuse pathology across neuronal processes and cell bodies could be linked to their higher tendency to fragment and diffuse in neurons. Interestingly, despite the structural and surface differences between the mouse and human fibril particles that include large differences in hydrophobicity, uptake remains the same in neurons through the same pathway.

### Mouse α-syn fibrils demonstrate exacerbated spreading in humanized α-syn mice

In order to explore whether the unique functional features of mouse α-syn fibrils discovered in primary neurons translate to the mouse brain, we selected SNCA-OVX transgenic mice that express human α-syn throughout their brain without expression of mouse *Snca*. We administered single intracranial injections into the dorsal striatum, using equal quantities of mouse or human PFFs and a vehicle control. Three months post-injection, a time when inclusions are described as most abundant (*13*, *86*, *87*), we stained the extracted brains against pS129-α-syn to investigate the spread and pathology abundance across the brain. We studied the propagation of α-syn pathology in each indicated brain region, comparing the ipsilateral and contralateral hemispheres (Fig. 6a). As anticipated, both PFF variants triggered the formation of pS129-α-syn in the ipsilateral hemisphere. The vehicle control injection, however, did not induce any α-syn-associated pathology, consistent with our past reports with this method (*88*). To further explore the transmission of α-syn pathology via inter-neuronal connectivity, we measured the spread of α-syn inclusions to the contralateral side. Our findings revealed that mouse α-syn PFF spread more effectively to the contralateral side in the piriform cortex, thalamus, and striatum (Fig. 6a-e). This result could not be explained by more pathological abundance at the injection site, since human PFFs caused more pathology in the ipsilateral dorsal striatum (i.e., around the injection site) than mouse PFFs (Fig. 5e).

**Fig. 6.**
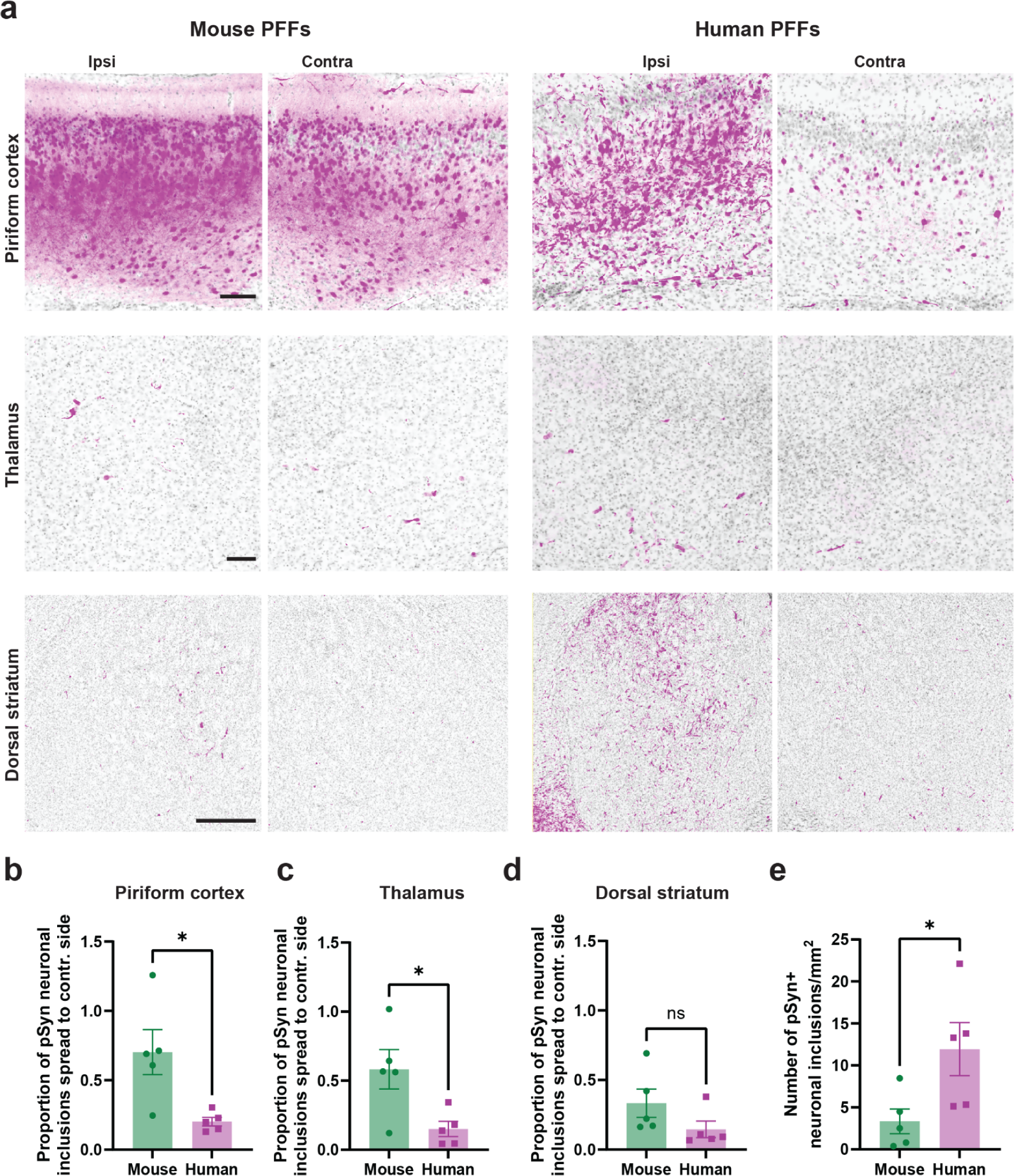
Progressive α-syn pathological spread differs between mouse and human α-syn PFFs. (a) Representative immunofluorescence images of pS129-α-syn pathology (magenta) and DAPI (grey) three months postinjection in human α-syn expressing SNCA-OVX mice unilaterally injected with mouse (left panel) or human (right panel) PFFs. Piriform cortex (top row), thalamus (middle row), dorsal striatum (bottom row) are separated to indicate ipsilateral (ipsi) and contralateral (contra) sides. Scale bars are 100 µm for top and middle rows and 500 µm for bottom row. Group analysis of α-syn pathology propagation ratio to contralateral side in piriform cortex (b), thalamus (c) and dorsal striatum (d) quantified as proportion of pSyn neuronal inclusion spread between ipsi and contra areas.(e) Quantitative analysis of pSyn neuronal inclusions per mm^2^ in striatum in mouse and human PFF cohorts. Each data point (n=5 per group) in group analysis plots represents the mean of the signal from 20-25 sections analyzed for each replicate and error bars indicating S.E.M. Significance was determined by 2-tailed t-tests; *p<0.05, ns for not significant

*In vitro* studies with mouse α-syn fibrils have demonstrated that they poorly template human α-syn protein into cross-seeded fibrils (*35*). Given the unexpected results here in both primary neurons and *in vivo* in a mouse model that expresses only human α-syn, we assessed again the cross-seeding properties of human and mouse α-syn fibrils using *in vitro* aggregation kinetics assays. Time-resolved aggregation assays were designed with normalized thioflavin T signals (i.e., percent of maximum signal) to account for the significant discrepancies in ThT binding between mouse and human fibrils. Consistent with previous observations, *in vitro* aggregation assays demonstrated that human PFFs are much more potent than mouse PFFs in templating human monomeric α-syn into new fibrils (Fig. 7a,b). Mouse PFFs poorly template human α-syn monomer, in contrast to what we observed in neurons. To confirm these results without the use of any amyloid dye, we repeated the experiments with endpoints defined by reactivity measured through the aggregation-selective MJFR-14-6-4-2 monoclonal antibody (Fig. 7c), which did not appear to have species preference between human and mouse. Using cross-labeled fibrils, microscopic inspection of the chimeric fibrils confirmed that both mouse and human PFFs are incorporated into single-growing fibrils harboring both mouse and human α-syn protein (Fig. 7d). Amyloid dye profiling of the chimeric products suggest that neither human nor mouse α-syn PFFs successfully preserve parental fibril structure into the newly born mixed (i.e., chimeric) fibrils. Instead, the cross-seeded products have intermediate amyloid dye profiles that are in-between the fibril parents (e.g., human to mouse, or mouse to human (Fig. 7e,f). Altogether, our findings are consistent with recent reports demonstrating notable differences in the seeding behavior *in vitro* compared to *in vivo*, even with the identical fibril preparation, and that the cellular environment as well as fibril mechanical properties plays critical roles in determining seeding potency and associated responses. Therefore, identifying specific cellular factors that drive *de novo* α-syn fibril formation and structure is essential to enable reproducing the structure of disease-associated brain-derived. On the basis of these observations, we recommend exercising caution in the potential over-interpretation of aggregation studies in cell-free systems.

**Fig. 7.**
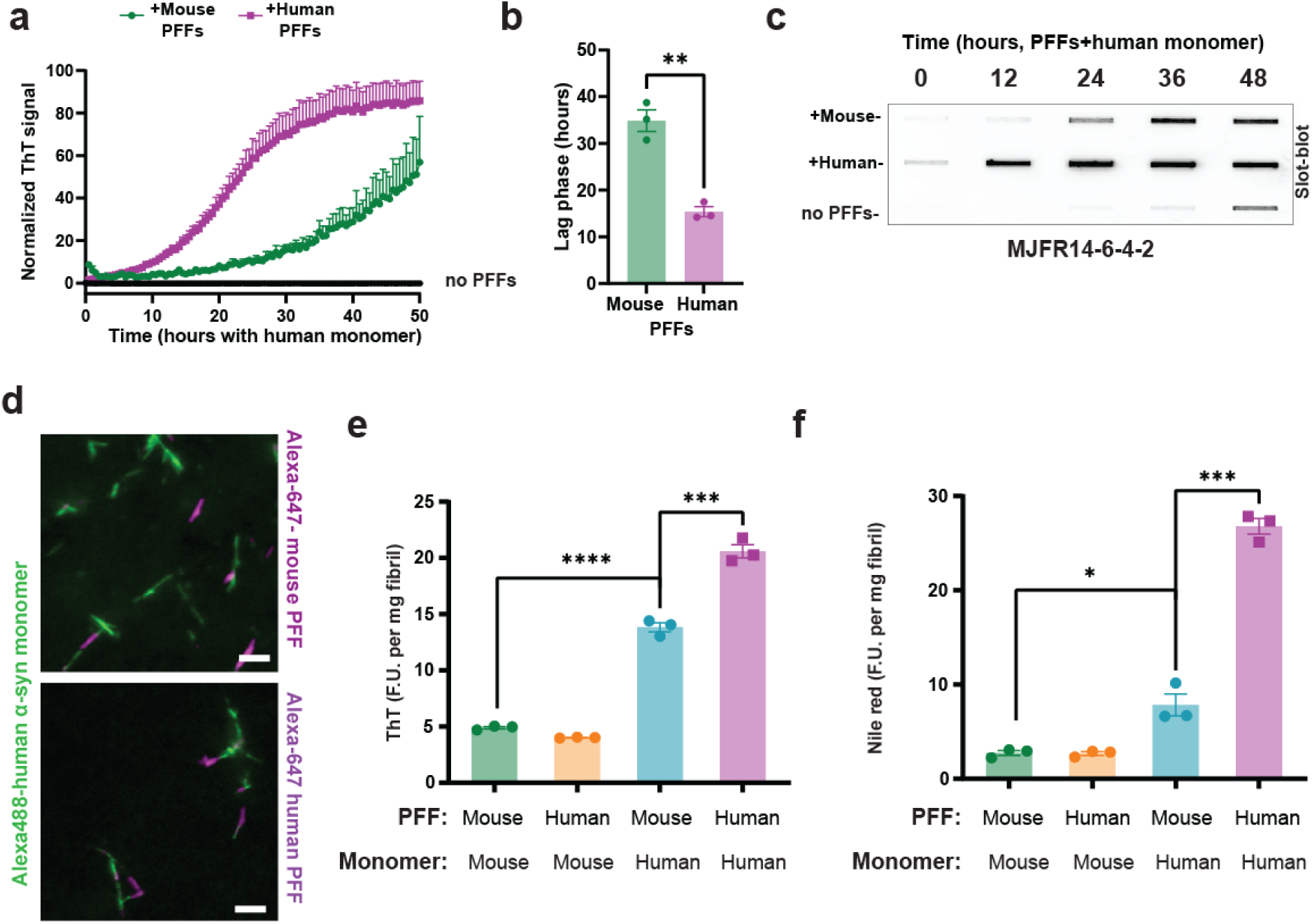
Recruitment of human α-syn monomer in mouse PFF-nucleated amplification leads to the formation of fibrils exhibiting distinct amyloid dye-binding profiles. (a) Representative RT-QuIC assay showing mouse and human PFF-templated aggregation with human α-syn monomer, including the indicated lag phase. Data points represent normalized ThT fluorescence from three independent experiments with error bars indicating S.E.M. (b) Group analysis of the duration of the lag phase in mouse and human PFF-templated aggregation. Dots represent a mean measured in duplicates from three independent experiments. (c) Representative filter-trap slot-blot analysis of dye-free aggregation of human α-syn monomer, with or without fibril PFFs, detected with MJFR14-6-4-2 α-syn aggregate-specific antibodies. (d) Representative images of Alexa-647-labeled mouse or human α-syn PFF-templated aggregation with human α-syn monomer, collected after 48 hours of incubation. The collected reactions were supplemented with 1 µM ThT (shown in green), with Alexa-647-conjugated PFFs depicted in magenta. The scale bar is 0.5 µm. Group analysis of ThT (e) and nile red (f) binding to extracted, monomer-free, sonicated α-syn fibrils (PFFs) performed in various combinations of mouse/human PFFs and their monomeric equivalents. Each data point in e and f represents a mean from an individual sample measured in duplicates from three independent experiments with error bars indicating S.E.M. *p<0.05, **p<0.01, **p<0.001, ****p < 0.0001 from 2-tailed t-tests.

## Discussion

Mouse α-syn fibrils are commonly used for the discovery of molecules that might bind to high affinity and function as probes for disease features, or in drug discovery efforts to find therapeutics that block aggregation (*89–91*). It has been presumed that the structures and features associated with mouse α-syn are close enough to those associated with disease that they will inform pathways and targets that might be pursued to better understand disease. Cryo-EM analysis of protein fibrils associated with disease offers unprecedented insights into how conformational variation in fibril structure might drive different disease endpoints. Despite the ubiquitous integration of mouse α-syn fibrils into research models of α-syn pathology formation and spreading in PD and other synucleinopathies, there have been no high-resolution structures of these tools to inform whether they are similar or distinct from the growing compendium of α-syn fibril conformations found in different diseases or generated under various aggregation conditions *in vitro*. Addressing this knowledge gap has significant implications for assessing the utility of seeding-based mouse models, which rely primarily on the induction of aggregation and pathology formation by the endogenous mouse α-syn, in translational PD research and drug discovery.

Herein, we report for the first time the cryo-EM structure of mouse α-syn fibrils and demonstrate that it exhibits distinct structural, amyloid-dyes binding, hydrophobicity profile, and stability properties compared to human α-syn fibrils prepared under identical conditions. Though different buffer conditions *in vitro* are known to affect fibril structures, fortunately, the field has largely coalesced around standardized *in vitro* growth conditions for mouse α-syn fibrils so that the structures and results produced in this study can be used to re-interpret past studies utilizing these particles. The high-resolution structure of mouse α-syn reveals a structural arrangement clearly distinct from human α-syn fibrils produced under the same buffer conditions. Moreover, the high similarity of mouse α-syn fibril cryo-EM maps independently generated and processed at two different sites, serves as a robust validation of the reproducibility inherent in our findings. The mouse α-syn fibril preparations were homogeneous and highly reproducible from one batch to the next, even between the two sites. Notably, the structure of the mouse fibrils does not resemble any of the brain-derived α-syn fibril structures from patients with PD and other synucleinopathies. Consistent with predictions from a comparative solid state NMR-based study on mouse and human α-syn fibrils, the interface between the protofilaments of mouse α-syn fibrils are more hydrophilic and the two types of fibril differ primarily at residues T53 (A53) and N87 (S87) which can broadly impact β6 and β7 strands within the amyloid core (*52*)immunogenic responses(*52*)mouse and human α-syn fibrils.

The deposition of protein fibrils into larger aggregates or their accumulation in intracellular inclusions has been associated with strong immunogenic responses in affected tissues of several neurodegenerative diseases (*75*, *92–96*). While a broad comparative analysis of immunogenic potential of different α-syn fibrils associated with disease has not been performed, to our knowledge, at least one study identified rod-like human α-syn fibrils as the most potent in inducing the secretion of pro-inflammatory cytokines and chemokines (*74*). The high expression of pattern recognition receptors like TLR4 are thought to drive these responses and the recognition of human α-syn fibrils at picomolar concentrations (*97*). In this work, we find that the more hydrophobic human 6sst α-syn fibrils are much more potent in eliciting an inflammatory response in macrophages, whereas mouse α-syn fibrils fail to cause a similar reaction. The Gal3 protein accumulates on impaired lysosomes and indicates the activation of autophagy-lysosomal pathways to eliminate hazardous molecules, including protein aggregates (*98–100*). The Gal3 protein acts as a TLR4 ligand to contribute to macrophage activation and IL-6 secretion (*101*). Though we did not pinpoint the mechanism of suppressed immune activation associated with mouse α-syn fibrils, it seems likely that the lack of lysosomal damage caused by the mouse fibrils prior to their elimination may cause a failure of toll-receptor signaling pathway and NLRP3 inflammasome activation that interconnects damaged lysosomes to the secretion of IL-6 and other cytokines in a variety of immune cells. Our findings suggest that the structural properties of the fibrils, and not only their biochemical properties, are key determinants of their interactions with lysosomes and pathogenicity (*87*).

While mouse PFF-induced inclusions in mice strongly correlate with dopaminergic neurodegeneration in non-transgenic mice (*13*, *46*), Our observation that mouse α-syn fibrils exhibit heightened fragmentation and increased sensitivity to detergents as well as increased spread of pathology in neurons may not be consistent with the general concept that fibril stability predicts pathogenicity (*58*). However, several studies have shown that the length of α-syn fibrils is a strong determinant of their seeding activity, with α-syn shorter fibrils exhibiting higher seeding activity (*46*, *62*, *63*). Fibril fragility in a cellular context, outside of the degradative lysosome, might be more advantageous for pathogenicity as enhanced fragmentation is thought to promote the spread of newly-seeded fibrils across the cell and across interconnected cells (*58*, *65*). However, what causes fibrils to fragment remains unknown, and it seems likely that factors in the cell milieu critically determine the pathogenicity of α-syn fibrils (*102*). A recent study by Ohgita et al investigated the relative contribution of intramolecular interactions between the C- and N-terminal domains of α-syn within fibrils on the stability of both mouse and human α-syn. Their results revealed that these intramolecular interactions are perturbed in mouse α-syn fibrils and could contribute to their reduced stability and higher tendency to fragment (*51*). Another study from our group showed that post-aggregation nitration induces α-syn fibril fragmentation *in vitro* (*103*). Sensitivity to denaturation or fragmentation has been postulated to be beneficial for pathogenicity in the disease (*58*), and mouse α-syn fibrils appear to closely mimic functional attributes described for MSA-associated fibrils. In support, MSA-tissue extracted α-syn fibrils show higher sensitivity to chemical degradation compared to PD-associated fibrils (*104*). Moreover, MSA fibrils are found to disassemble under lower concentrations of guanidine and exhibit higher pathology propagation potential compared to recombinant or PD-derived fibrils *in vitro* (*53*, *104*, *105*). Collectively, these results suggest other fibril properties associated with α-syn besides stability may be more important for α-syn-related pathogenicity and related disease phenotypes.

Our results underscore the critical importance of validating the structural similarities or differences of seeded aggregates to the native fibrils by cryo-EM before concluding successful amplification of native α-syn fibrils from brain or peripheral tissues. This is of crucial importance given the increasing reliance on *in vitro* amplification assays to draw conclusions about the structural and pathogenic properties of native α-syn fibrils. Comparison to other published α-syn fibril structures show clearly that mouse α-syn fibrils exhibit a high structural similarity to recombinant human E46K-mutated and human α-syn fibrils obtained by seeding human α-syn monomers with brain-derived fibrils from MSA brains, and that this similarity extends to functionality in seeding characteristics in neurons. Differential potency between human and mouse α-syn fibrils varies considerably from one neuron type to another, be it a human iPSC-derived dopaminergic neuron or a medium spiny neuron in the mouse dorsal striatum. It is clear through this study and others that both human and mouse α-syn PFFs associated with a variety of structures are available to cross- and potentially form new chimeric structures that do not resemble the parental structures. Compendiums that detail relative potencies and phenotypes associated with different fibril seeds may produce results relevant only to the structure and composition of the particular seed used in the selected model.

Following the results disclosed here, several predictions for the interpretation of past studies using mouse α-syn fibrils can be made based on the structure-to-function observations in this study. First, immunogenic responses to mouse α-syn fibrils may be underestimated compared to fibril types associated with Lewy body diseases. Second, the spread of pathology and induction of new fibrils templated from mouse α-syn fibrils in neurons may be overestimated compared to human fibrils, if not assessed using quantitative biochemical methods for monitoring loss of monomers or formation of fibrils. These findings may help explain why mouse α-syn fibrils are very successful in templating pathology compared to fibrils extracted from PD and DLB, regardless of whether the models endogenously express human or mouse *SNCA/Snca*. Third, *in vitro* aggregation assays, especially those exclusively reliant on thioflavin-binding, may not reflect important fibril properties that dictate pathological potency in cells that might include the effects of hydrophobicity and fragility differences between fibril types. Consistent with these observations and our findings, a recent study by Howe J et al reported that mouse α-syn fibrils induced more pS129 α-syn pathology in rat brains than rat α-syn fibrils, despite primary amino acid sequence differences, and suggested that additional cellular factors and not simply sequence homology between the fibrils and the monomeric subunits are the key determinants of α-syn fibrils seeding activity and pathogenic properties (*69*).

In summary, the results from this study have shed additional light on structural and functional properties of α-syn fibrils that may drive their pathobiological characteristics, and highlight the importance of a detailed structural analysis of tools used in research models that are intended to recapitulate central neuropathological features in certain aspects of disease. A better characterization of the types of α-syn aggregates disease, and methods to recapitulate them in models, may facilitate a higher success rate of pathological probes and therapeutics that mitigate the pathological effects of the aggregates in disease.

## Full Methods

### Purification of recombinant human and mouse α-synuclein

Recombinant expression and purification of human and mouse α-syn were conducted by a previously published protocol(*68*). Plasmid (pRK172) constructs encoding human and mouse α-syn proteins were used to transform *Escherichia Coli* B21 (DE3) codonplus cells (Agilent, Cat #230245-41). Protein expression at OD600 was induced by adding 0.05 mM IPTG (RPI, Cat #I56000) at 18°C for 16 hours. The collected bacterial pellets were thoroughly resuspended in lysis buffer (10 mM Tris-HCl, pH 7.6, 750 mM NaCl, 1mM PMSF, 1mM EDTA, protease inhibitors (Sigma-Aldrich, Cat #4693132001). Resuspended homogenates were lysed by sonication for 1 minute at 70% amplitude (Fisher500 Dismembrator, Fisher Inc.) and heating at 90°C for 15 minutes followed by centrifugation at 20k xg for 1 h at 4°C. Supernatants were dialyzed against the dialysis buffer (10 mM Tris-HCl, pH 7.6, 25 mM NaCL, 1 mM EDTA, 1mM PMSF) overnight and purified through HiPrep Q HP 16/10 column (Cytiva, Cat #29018182) with linear gradient application of high-salt buffer. Identified fractions were selected based on the presence of the protein band at correct molecular weight and the purity of the band evaluated by SDS-PAGE. Selected fractions were combined and then dialyzed with the dialysis buffer (10 mM Tris-HCl, pH 7.6, 25 mM NaCL) overnight. Anion exchange chromatography followed by dialysis was repeated to achieve a high purity sample. The dialyzed protein suspension was concentrated using ultracentrifugation units 3K cutoff (Amicon, Cat #UFC901024) up to 5 mL. Finally, endotoxins were removed using an endotoxin removal kit (Genscript, Cat #89233-330) and the levels of endotoxin were measured using an endotoxin detection kit (Genscript, Cat #95045-024). A detailed step-by-step protocol is provided at protocols.io dx.doi.org/10.17504/protocols.io.yxmvm3rz9l3p/v1

### Preparation of mouse and human α-synuclein fibrils

Monomeric α-syn at a concentration of 5 mg/mL in phosphate-buffered saline (PBS) was incubated for 5 days with intermittent shaking (1 minute on and off, Eppendorf, Thermomixer Model 5382, ThermoTop Smart Block Cat# 5308000003) at 1,000 xg. at 37°C. Resulted suspensions were centrifuged at 20k xg at 10°C for 10 minutes. Soluble fraction was removed and the remaining fibril pellets were washed three times in 1 mL of PBS. To measure fibril concentration, aliquots were titrated into 3 M guanidine chloride and agitated at room temperature for 5 minutes. Concentrations were then determined through A280 measurements using NanoDrop (Thermo Scientific, Cat# ND-ONE-W). In some experiments, fibril preparations were labeled with 100 μg of pHrodo iFL Red STP Ester (ThermoFisher, Cat# P36010), Alexa Fluor 647 NHS Ester (ThermoFisher, Cat# A20006), Alexa Fluor 568 NHS Ester (ThermoFisher, Cat# A20003), BODIPY-FL NHS ester (ThermoFisher Cat# D2184) at a concentration of 1 mg/mL in 1 mL of 100 mM sodium bicarbonate overnight with constant low-speed shaking at room temperature. Unconjugated dye was removed through rounds of pelleting the insoluble fractions via low-speed centrifugation and washing in PBS. For DQ fibrils equal concentrations of Alexa Fluor 647 and BODIPY-FL conjugated fibrils were mixed together prior to the cell treatment. Preceding the application as indicated, fibrils were sonicated to matched lengths of 20-30 nm in radius with polydispersity indices <0.5 determined by DLS conducted on the same day of the experiment. The sonication of fibrils was achieved with a cup horn sonicator in thin-walled 0.2 mL tubes (Q-Sonica, Cup horn, Cat #431C2; Q700 Sonicator Cat #Q700-110). The detailed protocol is described here: https://www.protocols.io/private/0B72006AD6B511EEBA4A0A58A9FEAC02

Fibrillization of chimeric fibrils was conducted by adding 1 % (50 µg/mL) recombinant mouse/human α-syn PFFs (or Alexa-647 conjugated variants) to the sample containing 5 mg/mL of mouse/human α-syn monomer in PBS. The reactions for the evaluation of fibril growth was incubated for 5 days with intermittent shaking (1 minute on and off, Eppendorf, Thermomixer Model 5382, ThermoTop Smart Block Cat# 5308000003) at 1,000 xg. at 37°C. To monitor the elongation of added PFFs by incorporation of monomeric α-syn, the resulting reactions were supplemented with 50 µM of ThT. After three wash cycles with PBS, the samples were further diluted to 400 ng/mL concentration and imaged on Zeiss Axio Observer Z1 using Plan-Apo oil immersion objective with 100x magnification.

### Cryo-electron microscopy and helical reconstructions

Full-length recombinant mouse and human α-syn fibrils were applied to plasma-cleaned (Pie Scientific) 1.2/1.3 UltrAufoil (Quantifoil, Cat #Q350AR13A) grids using an automatic grid plunger (Leica GP2) set at 95% humidity and at 20 °C. Prior to application to the grid, 4 µL aliquots of fibrils were thoroughly mixed. The mixed fibrils were then added to the grid with the blot parameters set for a duration of 4 seconds for back-blotting. Subsequently, the grids were plunge-frozen into a liquid Ethane/Propane mix. Images were captured on a Titan Krios cryo-TEM at a dose of 60 electrons per Å^2^, following the Latitude data collection procedures (Gatan, Inc.). Prior to atlas collection, image alignments and fine alignments were executed with selected data collection parameters outlined in Table 1. Movie frames were gain-corrected, aligned, and dose-weighed using Relion 4.0 software. CTF parameters were estimated using CTFFIND-4.1 software. For helical reconstructions, approximately 40 images were randomly selected for manual picking and around 14k fragments were extracted with an inter-particle distance of 6 β-rungs of 28.5 Å (with an estimated helical rise of 4.75 Å) from the start-end coordinate pairs. These extracted coordinates were used for training CRYOLO. The same model was used for all image datasets in auto picking mode with CRYOLO. Fragments were initially extracted using a box size of 1024 pixels and then down-scaled to 256 pixels, resulting in a pixel size of 4.32 Å. Reference-free 2D classifications were performed to assess different fibril strains, cross-over distances (estimated as 1,200 Å), and helical rise (4.82 Å), and to select suitable fragments for further processing. Initial references were generated from selected 2D class averages using the Relion_helix_inimodel2d command with a cross-over distance of 1200 Å, search shift of 15 Å, and search angle of 15 degrees. The 2D classes showing splitted protofilaments or protofilaments only were discarded. Another round of 2D classification was performed using a box size of 384 and no binning. The clean set of fragments achieved by selecting 2D classes showing clear rungs features was re-extracted using box sizes of 512 pixels and down-scaled to 256 pixels (binned pixel size of 2.16 Å) for optimal performance in 3D classifications. To minimize reference bias, initial models were low-pass filtered to 40 Å. During 3D classification, 6-8 classes were used along with initial helical twists set to −0.7 degrees, initial helical rises of 4.82 Å, and a regularization parameter T of 4. Helical symmetry local search was performed during classification with helical twist ranges varying from −0.85 to −0.5 and helical rise ranges varying from 4.8 to 4.84. Only the central 10% of the z length was used for reconstructions. Classes with nominal folding features were individually selected and re-extracted with box sizes of 384 pixels. Additional rounds of 3D classification were performed with the new reconstructions as reference using an initial low-pass filter set to 10 Å to further clean the particles. Initial helical twist and helical rise parameters were updated from 3D classifications. Local symmetry search was not performed at the first round of refinement. In the second round refinement, a shape mask comprising 90% of the central z length was applied along with local symmetry search. To improve the resolution, multiple rounds of CTF refinement, Bayesian polishing, and refinements with previous results using a particle extraction size of 256 pixels were used. Post processing was accomplished using a shape mask comprising 10% of the central z length, and final resolutions were estimated for each dataset using the gold standard FSC 0.143 cutoff. The final twist and rise for mouse fibril were determined as +/-0.84 and 4.84, respectively. And the final twist and rise for human spontaneous fibril were determined as +/-0.75° and 4.82 Å, respectively. More information about the processing is included in Table S1. The detailed protocol is described here: https://www.protocols.io/private/7A7AAF9DDBFD11EEBBF10A58A9FEAC02

#### Cryo-EM helical image processing and model refinement (EPFL site)

Cryo-EM images were collected using a Thermo Fisher Scientific (TFS) Glacios electron microscope, equipped with an X-FEG electron source and operated at 200kV. A total of 5,258 dose-fractionated movies were recorded with a Gatan K3 camera in counting mode at a physical pixel size of 0.8780 Å, a defocus ranging from 0.8 to 3 µm, and a total dose of ∼50 e-/Å2, using SerialEM with life analysis using FOCUS (*106*, *107*). The movies were imported into RELION 4.0, for data processing (*108*, *109*). Motion correction was performed using RELION’s implementation of Motioncor2 with a dose of 1.6 e-/Å2 per frame(*110*). CTF correction was done using CTFFIND 4.1.14, and 4,904 selected micrographs were used for further image processing (111). From the 4,904 micrographs with a maximum CTF resolution better than 5 Å and defocus ranging from 0.8 to 2.4 µm, 4,144 fibrils were manually selected. At first, 271,844 particles with a box size of 674 Å were extracted and subjected to reference-free 2D classification, leading to the selection of 146,075 particles, which were subsequently re-extracted with a box size of 316 Å (360 pixels) for 3D refinement. The initial model was created using IniModel2D from six distinct 2D averages, each having a box size of 674 Å, encompassing diverse views and a crossover distance of 1050 Å. After two rounds of 3D refinement, a third refinement was tested for C2 and pseudo-2-fold symmetry. The structure with pseudo-2-fold symmetry, was implemented in 3D refinement and classifications. After multiple rounds of 3D classification and refinement (helical z-percentages between 0.15-0.3), a total of 50,378 particles with 316 Å box size underwent CTF refinement, Bayesian polishing, and post-processing, yielding an optimized twist 179.598° and a rise of 2.39 Å. Post-processing was performed with a soft-edged mask and an estimated sharpening B-factor of −52 Å2, resulting in a 2.9 Å structure according to gold-standard FSC - 0.143 criteria. Model building was performed manually for one rung of the single protofilament, using Coot 0.9.8.92(*112*). Both unsharpened and sharpened maps were used for modeling the peptide backbone. Further, rounds of iterative real-space refinement were carried out on six adjacent fibril stackings of the sharpened map using PHENIX (phenix.real_space_refine), and ISOLDE, incorporating β-stacking secondary structure strains(*113*, *114*)(*115*). The model quality was assessed using Molprobity (*116*). The matchmaker function in ChimeraX was employed to facilitate comparisons with previously published structures(*117*).

### Structure alignments

Mouse α-syn structure (8uie) was superimposed and aligned with recombinant human (6sst), recombinant E46K (6ufr), and MSA-amplified (7ncg) in UCSF ChimeraX using the command “align” focusing on residue range 54-66 where the 3D feature indicated the most conserved among those structures(*118*). Cα RMSDs values were calculated based on the alignment with sequence range of 34-97. For global alignment mouse α-syn fibrils (8uie), recombinant human (6sst), recombinant E46K (6ufr), and MSA-amplified (7ncg) were imported into Maestro (Schrodinger Inc.) and prepared with the Protein Preparation Workflow task. Preprocessing was performed with the following parameters: fill in missing side chains; assign bond orders, using Combined Chemical Dictionary (CCD) database; replace hydrogens; create zero-order bonds to metals; create disulfide bonds; fill in missing loops using Prime; sample water orientations, use crystal symmetry, minimize hydrogens of altered species, use PROPKA with pH 7.4; restrained minimization was then performed, converging heavy atoms to root mean square deviation of 0.3 Å using OPLS4; waters were removed 3.0 Å beyond het groups. The prepared fibril structures were trimmed on their terminal ends to each contain five β-structure repeat units and amino acids 37-96 (600 residues total for each fibril). A single protofilament was isolated by removal of the paired protofilament in the fibril structure. Prepared protofilament structures were exported to PDB format for alignment. Protofilament structures were aligned using MM-align and RMSD values were computed(*119*). MM-align computes alignments of multi-chain protein complexes. Aligned structures were exported from MM-align to PDB format.

### MD simulations

For the preparation of starting configurations of molecular dynamics trajectories, which were based on the cryoEM structures, missing atoms and protons were introduced by using the leap module of Amber.20. Counter ions were added, and the systems were solvated in a box of water with the box boundary extending to 20 Å from the nearest peptide atom. Prior to equilibration, the solvated system was sequentially subjected to 1) 500 ps belly dynamics with fixed peptide, 2) minimization, 3) low temperature constant pressure dynamics at fixed protein to assure a reasonable starting density, 4) minimization, 5) stepwise heating MD at constant volume, and 6) constant volume simulation for 10 ns with a constraint force constant of 10 kcal/mol applied only on backbone heavy atoms. After releasing all constraining forces, sampling was increased by performing 4 independent, constant-temperature constant-volume molecular dynamics simulations for 100 ns each. All trajectories were calculated using the PMEMD module of Amber.20 with 1 fs time step. The amino acid parameters were selected from the SB14ff force field of Amber.20 and configurations were applied for every ns of the analysis.

### Tensile strength MMGBSA energy analysis

Using the Molecular Mechanics with Generalized Born and Surface Area (MMGBSA) protocol of Amber.20, interaction free energies of all the residues in the 3rd protofilament with the residues in the two surrounding protofilament from each side were calculated for the 100 configurations extracted at each ns interval from each trajectory. The ionic strength for MMGBSA calculations was selected to be 0.1M.

### Amyloid dye binding assays

Fresh aliquots of 50 mM nile red (Sigma, Cat #19123-10MG), FSB (Sigma, Cat #344101-5MG) and 500 mM ThT (Sigma-Aldrich, Cat #BC85) in DMSO were diluted in PBS to reach 100 µM concentration. Freshly sonicated α-syn PFFs evaluated by DLS were prepared at concentration 1 mg/mL followed by serial dilutions by a factor of 2. Diluted α-syn PFFs were combined with amyloid dye aliquots and transferred into 384-well plates with clear bottoms (Corning, Cat #4588). Fluorescence intensity was recorded on CLARIOstar OMEGA reader (BMG, Inc.) with excitation/emission spectra set at 468-15/510-20 for FBS, 535-20/585-30 for nile red and 448-10/482-10 nm for ThT. Monomeric mouse and human α-syn monomers were used as a control. A detailed step-by-step protocol is provided at protocols.io: https://www.protocols.io/private/55CB49A4E35A11ECB9FC0A58A9FEAC02

For real-time aggregation reactions, samples were supplemented with 10 μM ThT in ultra-low binding 384-well plates with clear bottoms and sealed with foil (BioRad, Cat #1814040). Each plate was supplemented with a standard curve of serial dilutions of recombinant α-syn reference PFFs. Reaction fluorescence was monitored at 448-10 nm excitation and 482-10 nm emission on FLUOstar Omega (BMG, Inc.) plate reader every 30 minutes with intermittent shaking at 700xg. A detailed step-by-step protocol is provided at protocols.io: dx.doi.org/10.17504/protocols.io.6qpvr67kpvmk/v1

### Fibril fragmentation and denaturation analyses

Freshly prepared full-length α-syn fibrils were diluted to 5 mg/mL and aliquoted into 0.2 mL thin-wall PCR tubes (BrandTech, Cat #13-882-58), each containing 20 µL of the sample. These aliquoted α-syn fibrils were then placed into a PCR tube adaptor (Qsonica, Cat #451) and subjected to ultrasonic waves using a cup horn water-bath sonicator (QSonica, Cat #431C2). The sonication was performed for 2 minutes at 30% amplitude in a temperature-controlled environment set at 10°C. After sonication, the samples were collected for subsequent analysis. The sonicated fibrils were further diluted to 200 ng/mL and measured using the DLS approach, conducting 30 acquisitions for each sample. A separate set of freshly prepared full-length α-syn fibrils underwent sonication for 30 minutes. For the analysis, size distributions were divided into groups of sizes: 1-10 nm, 10-100 nm, >100 nm. Each sonication cycle included three technical replicates to account for the variability of the distribution of ultrasonic waves in the cup horn adaptor. A detailed step-by-step protocol is provided at protocols.io: https://www.protocols.io/private/ACD8D8CCD66311EEBBE90A58A9FEAC02

For the chemical denaturation analysis, sonicated fibrils were initially assessed using DLS to ensure that the size distributions matched between the mouse and human fibril preparations. Chemical denaturants were subjected to serial dilutions, with a dilution factor of 10 for SDS (Sigma-Aldrich, Cat #436143) starting from 10%, 2 for sarkosyl (N-Lauroylsarcosine, Sigma-Aldrich, Cat #61739) starting from 2%, and GuHCL (Sigma-Aldrich, Cat #G3272) starting from 6M concentrations. The sonicated fibrils were then diluted to a concentration of 2 mg/mL and spiked into the serial dilutions of the chemical denaturants. These reactions were incubated for 30 minutes and then diluted to 200 ng/mL prior to the slot-blot analysis. A detailed step-by-step protocol is provided at protocols.io: https://www.protocols.io/private/564D3906DCAE11EE9AD80A58A9FEAC02

### Slot-blot analysis

Samples containing 200 µL of α-syn PFFs at 200 ng/mL concentration were applied to previously soaked in TBS 0.22 µm nitrocellulose membrane using a bio-dot slot format microfiltration apparatus (BioRad, Cat #1706542). Samples were filtered and washed in TBST through the membrane using a low-speed vacuum to ensure full drainage of the liquid. Washed membranes were removed from the cassette and placed in the blocking solution containing 5% w/v nonfat dry milk (Bio Rad, Cat #1706404XTU) in TBST. The membrane then was transferred into a solution containing Alexa-647-MJFR14-6-4-2 (Abcam, Cat #ab216309) antibodies for detection of α-syn-specific aggregates. A detailed step-by-step protocol is provided at protocols.io: https://www.protocols.io/private/8A5B8C8DDCAF11EE9AD80A58A9FEAC02

### MDM culture

Mononuclear cells from the blood of healthy volunteers were separated using SepMate and Lymphoprep tubes (Stemcell Tech, Inc.), as previously described(*76*). The cells were then processed using EasySep Negative Selection Human Monocyte Enrichment kits (Stemcell Tech, Cat #19058), without depleting CD16 and then cultured in DMEM (Invitrogen, Cat #11995073) supplemented with Glutamax (ThermoFisher, Cat #35050061), 10% fetal bovine serum (FBS; Atlanta Biologicals, Cat #S11150H), Antibiotic-antimycotic (ThermoFisher, Cat #15240062), Animal-Free Recombinant Human M-CSF (20 ng/mL, PeproTech, Cat#AF-300-25). The cells were cultured for 7 days before the experiments to obtain human macrophage characteristics and treated with α-syn PFFs. MDM cultures incubated with α-syn PFFs for periods of time indicated in figure legends then washed thrice with endotoxin-free PBS from (Sigma-Aldrich, Cat #TMS-012-A). After three wash cycles with endotoxin-free PBS on ice, MDM cells were fixed with 4% PFA (EMS, Cat #15700). PBS supplemented with 5% BSA (Sigma-Aldrich, Cat #A4612), FITC pre-conjugated LAMP1 Monoclonal Antibody (LY1C6; 1:100, Thermo Fisher Scientific, Cat #MA1-164, RRID AB_2536869) or Galectin 3 Monoclonal Antibody (A3A12; Thermo Fisher Scientific, Cat #MA1-940, RRID AB_2136775), and Hoechst 33342 (1:5000, BD Bioscience, Cat #561908, RRID AB_2869394). A detailed step-by-step protocol is provided at protocols.io: https://www.protocols.io/private/1D2E7303EC0C11ECB2DC0A58A9FEAC02

### Primary neuron cultures preparation and analysis

All procedures involving mice were approved by the Duke Institutional Animal Care and Use committee. Primary hippocampal neuronal cultures were prepared from postnatal (P0) pups of nTg (The Jackson Laboratory, JAX stock #000664, RRID IMSR_JAX:000664), human-PAC-wt-*SNCA*^+/+^/*Snca*^−/−^(JAX stock #010710, RRID IMSR_JAX:010710) or *Snca*^−/−^(JAX stock #016123, RRID IMSR_JAX:016123) and cultured as previously described(*68*). Briefly, the hippocampi were isolated and dissected in Hibernate E medium (VWR, Cat #MSPP-HE). The tissue was then digested in a papain solution (Worthington, Cat #LS003126) in an HBSS buffer (Thermo Fisher Scientific, Cat #14025126), supplemented with 10 mM HEPES pH 7.4, 100 mM sodium pyruvate, and 1% penicillin/streptomycin (Thermo Fisher Scientific, Cat #10378016). The cells were plated and incubated in Neurobasal media (Thermo Fisher Scientific, Cat #21103049) with B-27 (Thermo Fisher Scientific, Cat #17504044), 5mM GlutaMAX, and 10% FBS (Atlanta Biologicals, Cat #S11150H) in wells coated with poly-d-lysine (0.1 mg/ml; Thermo Fisher Scientific, Cat #A3890401) at a density of 10,000 cells per cm^2^. After 12 hours, the plating media was replaced with a maintenance medium that included a Neurobasal medium supplemented with B-27 and 0.5 mM L-glutamine. The primary cultures that reached DIV7 were used in experiments. A detailed step-by-step protocol is provided at protocols.io: https://www.protocols.io/private/E4AE6DC5DFB511EC88D20A58A9FEAC02

Primary hippocampal neurons at DIV7 prepared according to the aforementioned protocol were treated with α-syn PFFs at concentration 0.64 nM or 64 pM (as indicated in figure legends as 10 µg/mL or 1 µg/mL) relative to estimated molecular weight measured as calculated size by DLS acquisitions and protein weight measured via Nanodrop. For ELISA measurements, primary neuron cultures in 48-well plates at DIV21 were lysed in 200 µL of a lysis buffer containing 1% Triton, protease and phosphatase inhibitors in PBS. Cells were thoroughly scraped and resuspended before transferring into eppendorf tubes. The collected samples were then sonicated using a water-bath sonicator (QSonica700, QSonica Inc.) at 30% amplitude at 10°C for a duration of 10 minutes. The sonicated cell suspensions were then centrifuged at 10k xg for 20 minutes at 4°C. The supernatants were aliquoted and stored at +/-80°C for subsequent BCA analysis and ELISA measurements. For immunofluorescence analysis, culture media at DIV21 was removed and fixed with 2% PFA (EMS, Cat #15700) in TBS. Samples were then permeabilized in 3% normal donkey serum (Equitech-Bio, Cat #SD32-0500) supplemented with 0.1% saponin (Sigma-Aldrich, Cat #S4521-25G) in TBS for 30 minutes. Primary antibodies including anti-α-syn (phospho S129) antibody EP1536Y (1:4,000, Abcam, Cat #ab51253, RRID AB_869973), Tau5 (1:2,000, Thermo Fisher Scientific, Cat #AHB0042, RRID AB_1502093), NeuN (1:2,000, GeneRwx, Cat #GTX00837, RRID AB_2937041) were incubated in 0.02%Saponin, 1% normal donkey serum in TBS overnight at 4°C with gentle agitation. After three constitutive washing rounds with TBS secondary antibodies including anti-rabbit Alexa Fluor 647 (Thermo Fisher Scientific, Cat #A-31573, RRID AB_2536183) anti-mouse Alexa Fluor 488 (Thermo Fisher Scientific, Cat #A-21131, RRID AB_2535771), anti-chicken Alexa Fluor 555 (Jackson Immuno, Cat #703545155, RRID AB_2340375), Hoechst 33342 (1:10,000) overnight at 4°C with orbital shaking. Images were obtained using Keyence BZ-X810 and Zeiss880 and coded for a blinded approach to analyze using CellProfiler and imageJ. A detailed step-by-step protocol is provided at protocols.io: https://www.protocols.io/private/BA28A193DCB311EEAE260A58A9FEAC02

### iPSC DA preparation and analysis

The iPSCs were differentiated into midbrain dopaminergic neurons (DA neurons) following the previously published protocol (*120*, *121*). Neurons were cultured in Neurobasal SM1 media (Thermo Fisher Scientific, Cat #21103-049) containing NeuroCult SM1 supplement (StemCell Technologies, Cat #05711), 1% penicillin / streptomycin (Thermo Fisher Scientific, Cat #10378016), and 1% L-glutamine (Gibco, Cat #25030081). Neurons were aged to 60-70 days for each experiment as indicated. Neurons were cultured on either 48-well (8 mm; Warner Instruments, Cat #64-0701) or 24-well (12 mm; VWR, Cat #48366-251) sized coverslips sequentially coated with poly-d-lysine (Sigma Aldrich, Cat #P1149-10mg) and laminin (Sigma Aldrich, Cat #11243217001). Before undergoing differentiation, all iPSCs were maintained in mTeSR1 media (StemCell Technologies, Cat #85850) on Cultrex (Thermo Fisher Scientific, Cat #343301001) coated plates. For iPSC DA neuron characterization, the 60–75-day old cultures were permeabilized and blocked with a blocking solution comprised of 0.1% Triton X-100 in PBS (Thermo Fisher Scientific, Cat #10010-049) supplemented with 2% BSA (Roche, Cat #03117057001) and 5% normal goat serum (Jackson ImmunoResearch, Cat #005-000-121) overnight at 4°C to prevent non-specific antibody binding. Neurons were incubated with primary antibodies against β-III-tubulin (1:333, Biolegend, Cat #802001, RRID AB_2564645), Tyrosine hydroxylase (1:333, TH; EMD Millipore, Cat #AB9702, RRID AB_570923), FOXA2/HNF3β (1:333, Santa Cruz, Cat #sc-101060, RRID AB_1124660) in blocking solution overnight at 4°C, washed three times with a 0.1% Triton X-100 PBS wash solution, followed by a 1 hour, room temperature incubation with secondary antibodies Alexa Fluor 568 Donkey anti-Rabbit IgG (Invitrogen, Cat #A10042, RRID AB_2534017), Alexa Fluor 488 Goat anti-Chicken IgY (Invitrogen, Cat #A32931, RRID AB_2762843), and Alexa Fluor 647 Donkey anti-Mouse IgG (Invitrogen, Cat #A-31571, RRID AB_162542) all at 1:500. The cells were then washed three times with wash solution and mounted onto microscope slides (Fisher Scientific, Cat #12-550-15) with DAPI Fluoromount mounting media (Southern Biotech, Cat #0100-20) and imaged. For seeding experiments 60–75-day old iPSC DA neurons cultured on coverslips were treated with Alexa 568 mouse and human α-syn PFFs at a concentration of 10 μg/ml. During the seven days of incubation, the media was changed every 48 hours. DA neurons were immunostained as described above with β-III-tubulin(1:500, Abcam, Cat #ab41489, RRID AB_727049), anti-α-syn (phospho S129) antibody EP1536Y (1:2,000, Abcam, Cat #ab51253, RRID AB_869973) and DAPI Fluoromount-G (Southern Biotech, Cat #0100-20).

### Endocytosis and internalization assays

For endocytosis assay in primary hippocampal culture curated from nTg (The Jackson Laboratory, JAX stock #000664, RRID IMSR_JAX:000664) P0 pups at DIV7 were treated with endocytosis inhibitors including 50 μM ethyl-isopropyl amiloride (EIPA, Caymanchem, Cat #14406), 50 μM Dyngo 4a (Abcam, ab120689), 0.2 μM wortmannin (Sigma-Aldrich, Cat #W1628-1MG), 15 μM Pitstop 2 (Abcam,Cat #ab120687), or 2 mM methyl-β-cyclodextrin (Sigma, Cat #C4555-1G) or 0.001% dimethyl sulfoxide (DMSO, Millipore, Cat #D2650-5X5ML; vehicle control) for 30 minutes before incubation with pHrodo-conjugated α-syn PFFs. The internalization rate was measured by the proportion of fluorescence emitting from the conjugated fibrils and DAPI count with added cell mask. A detailed step-by-step protocol is provided at protocols.io: https://www.protocols.io/private/E83B2912DCB311EEAE260A58A9FEAC02

Uptake of ∼60-day old iPSC DA neurons were assessed by the treatment of Alexa 568 human and mouse α-syn PFFs at 10 μg/ml compared to untreated control cells to measure uptake. Following 8 hours of treatment, DA neurons were fixed in 4% PFA (Polysciences, Cat #40181) for 30 minutes then washed with PBS (Thermo Fisher Scientific, Cat #10010-049) three times. The total cell membrane was labeled using CellTag 700 (Li-Cor Biosciences, Cat #926-41090) according to the manufacturer’s instructions. In brief, CellTag 700 was reconstituted in 100 μl of PBS, vortexed for ∼45 seconds, and allowed to rehydrate in the dark for 30 minutes. The reconstituted CellTag 700 was diluted 1:500 and 300 μl and added to a coverslip within a 48-well plate. Following a 1-hour room temperature incubation protected from light, the cells were washed 3x with PBS, and 300 μl of 0.1% Trypan Blue (Invitrogen, Cat #T10282) was added for 5 minutes to quench extracellular α-syn fibril signal. The wells were aspirated, and the coverslips were mounted onto microscope slides (Fisher Scientific, Cat #12-550-15) with DAPI Fluoromount mounting media (Southern Biotech, Cat #0100-20) and imaged.

For internalization analysis in MDM cultures, cells were treated with Alexa 647 human and mouse α-syn PFFs at 1 μg/ml for the indicated time followed by subsequent wash cycles with PBS, fixation in 4% PFA in PBS and immunostained for LAMP1 and Hoechst 33342.

### Enzyme-linked immunosorbent assays (ELISA)

Human and mouse α-syn aggregate levels was measured according to the manufacturer’s protocol (Biolegend, Cat #449407). In brief, collected cell lysates were diluted (1:100) prior to adding to the antibody-coated wells along with two separate standard curves of serially diluted mouse and human α-syn PFFs. Absorbance was recorded at 450 nm using SPECTROstar plate reader (BMG, Inc.).

A detailed step-by-step protocol is provided: https://www.protocols.io/private/055FBA22DA4611EC90060A58A9FEAC02 Chemokines and cytokines were analyzed via R&D DuoSet ELISA systems (Human CCL5/RANTES DuoSet ELISA, Cat #DY278; DuoSet ELISA Ancillary Reagent Kit 2, Cat #DY008; Human IL-6 Elisa Kit, Cat #DY206-05; R&D systems) according to manufacturer’s instructions and include human IL-6 and CCL5.

### Stereotaxic intracranial injections and processing

Intracranial injections of α-syn fibrils, or vehicle control, were performed as previously described (*46*). Transgenic mice SNCA-OVX (B6.Cg-Tg(SNCA)OVX37Rwm Snca^tm1Rosl^/J) obtained from Jackson Laboratories (JAX stock #023837; RRID:IMSR_JAX:023837 were randomized to groups spread across multiple cages. Stereotaxic injections of α-syn PFFs (10 μg per injection, equivalent protein weight for both human and mouse variants), were performed on mice at 4-5 months of age. A solution containing the indicated amount of α-syn PFFs (2 μl per each injection), or vehicle control, was injected into the right dorsal striatum relative to bregma: 1.0 mm anterior, 1.85 mm lateral, and 3.0 mm ventral relative to the skull. Three months after injections, mice were anesthetized with isoflurane and transcardially perfused with PBS followed by freshly prepared 4% PFA buffered in PBS. Removed brains were postfixed for 12 hours in a 4% PFA in PBS followed by an incubation in a 30% sucrose PBS for 48 hours. Frozen brains were then sectioned to 40 μm using a freezing microtome (Leica SM2010 R Sliding Microtome) and placed into 6-well plates with 50% glycerol in PBS. Sections were incubated in an antigen retrieval buffer (10 mM sodium citrate, pH 6.0, supplemented with 0.05% Tween 20) for 30 min with gentle rocking at 37°C followed by three wash cycles in tris-buffered saline (TBS; Thermo Fisher Scientific, Cat# J60764.K2). Rinsed sections were incubated in 5% donkey serum and 0.3% Triton X-100 in TBS for 1 hour at RT. For α-syn pathology detection, sections were incubated with anti-α-syn (phospho S129) antibody EP1536Y (1:2500; Abcam, Cat #ab51253, RRID AB_869973) primary antibodies in TBS buffer supplemented with 2.5% donkey serum and 0.1% Triton X-100 for 24 hours. Followed by wash cycles in TBS, sections were then incubated with donkey anti-rabbit Alexa Fluor 488 (1:500; Abcam, Cat #A32790, RRID:AB_2762833) secondary antibodies and Hoechst 33342 (1:5000, BD Bioscience, Cat #561908, RRID AB_2869394) for 24 hours. Immunostained sections were mounted on superfrost slides with ProLong Gold Antifade Mountant (Thermo Fisher Scientific, Cat# P36930) and stored at 4°C until imaging on the slide scanner VS200 Olympus. To quantitatively measure the distribution of pSyn-positive areas, an AI model designed in QuPath software was implemented and applied to each collected brain section image. Further sections were imported into QuickN software to align with the Allen Brain Atlas mouse brain edition (2017) followed by further adjustment using Visualign software.

pSyn area calculations were performed using Nutil software and data was analyzed as the proportion of pSyn area to the brain region area for each section. The average of 15-25 collected sections per animal was calculated to generate a data point used to build a group analysis plot in GraphPad software. Protocol used as a reference is detailed here: dx.doi.org/10.17504/protocols.io.4r3l22y6jl1y/v1

Analysis was performed by investigators blinded to sample identity until final data curation using coded identifiers for slides, mice and injected material.

### α-Synuclein sedimentation

Seeding and fibrillization growth was assayed by adding 1% α-syn PFFs (3 μg/mL) into a fibrillization reaction that contained 0.3 mg/mL of freshly thawed mouse or human α-syn monomer. The fibrillization reactions were performed according to the RT-QuIC approach amyloid dye-free environment in triplicate, with a volume of 50 μL for each specified time point. The resulting reactions were analyzed using the previously described slot-blot analysis or through SDS-PAGE with a coomassie stain. A detailed step-by-step protocol is provided at protocols.io:

dx.doi.org/10.17504/protocols.io.6qpvr3nwpvmk/v1

### Statistical analyses

Statistical analyses were performed using the GraphPad Prism 9 software. The specific statistical tests used for each dataset are detailed in the legends of each figure.

## Acknowledgments

We thank Lisa Cameron and Duke Light Microscopy Core Facility for support of fluorescence microscopy image collection and analysis, Nilakshee Bhattacharya and Duke SMIF cryo-EM core for assistance with and maintenance with cryo-EM collection, and Duke University Research Computing for maintenance and providing use of the Duke Compute Cluster (DCC). Michael Henderson for support in adapting QUINT workflow for brain image analysis. We thank the BioEM Lab of the University of Basel for the cryo-EM data collection of the EPFL structure.

## Funding

This research was funded in whole or in part by Aligning Science Across Parkinson’s Grand ID #020527 through the Michael J. Fox Foundation for Parkinson’s Research (MJFF). MJFF administers the grant ASAP-020527 on behalf of ASAP and itself. This work was also supported by the Swiss National Science Foundation, grant 310030_188548.

## Data Availability

Raw micrographs are available at the Electron Microscopy Public Image Archive, accession number EMPIAR-*In progress*. The reconstructed 3D maps are available at the Electron Microscopy Data Base, accession number EMD-50023 and EMD-42294. The atomic map is available at the Protein Data Base, accession number PDB-9EWV and 8UIE

## Supplemental figures

**Table S1.** Statistics of cryo-EM data collection and structure refinement of human α-syn fibrils.

| **Fibril type:** | **Human (similar to 6sst)** | **Human (6sst)** |
| --- | --- | --- |
|  |  | adapted from (21) |
| **Data collection** |  |  |
| Pixel size (Å) | 1.08 | 0.629 |
| Defocus Range (μm) | -2.5 to -0.8 | -0.8 to -2.5 |
| Voltage (kV) | 300 | 300 |
| Exposure time (s/ movie) | 60? | 10? |
| Number of frames | 55 | 50 |
| Total dose (e<sup>-</sup>/ Å<sup>2</sup>) | 55 | 69 |
| Micrographs | 6,530 | 1,143 |
| Inter-box distance (pixels) | 27 | 28 |
| Initially extracted fragments | 1,689,529 | 100,323 |
| Segments after 2D classification | 526,795 | 100,193 |
| **Reconstruction** |  |  |
| Segments after 3D classification | 81,190 | 3,989 |
| Resolution after 3D refinement (Å) | 2.8 | 3.75 |
| Final resolution (Å) | 2.7 | 3.39 |
| Estimated map sharpening B-factor (Å<sup>2</sup>) | -68 | -76.4 |
| Helical rise (Å) | 4.82 | 2.40 |
| Helical twist (°) | -0.75 | 179.55 |
| **Atomic model** |  |  |
| EMDB code | N/A | EMD-10305 |
| PDB code | N/A | 6sst |
| FSC threshold | 0.143 | 0.143 |
| **Composition** | N.D. |  |
| - Chains |  |  |
| - Atoms(non-Hydrogen) |  | 4900 |
| - Residues |  | 730 |
| **B-factors (Å<sup>2</sup>)** |  |  |
| -Protein |  | 70.32 |
| **R.M.S. deviations** |  |  |
| - Bond lengths (Å) |  | 0.009 |
| - Bond angles (°) |  | 0.941 |
| **Validation** |  |  |
| - MolProbity score |  | 2.11 |
| - Clashscore |  | 5.73 |
| - Poor rotamers (%) |  | 2.13 |
| **Ramachandran plot** |  |  |
| -Favored (%) |  | 89.86 |
| -Allowed (%) |  | 10.14 |
| -Disallowed (%) |  | 0.00 |

**Table S2.**
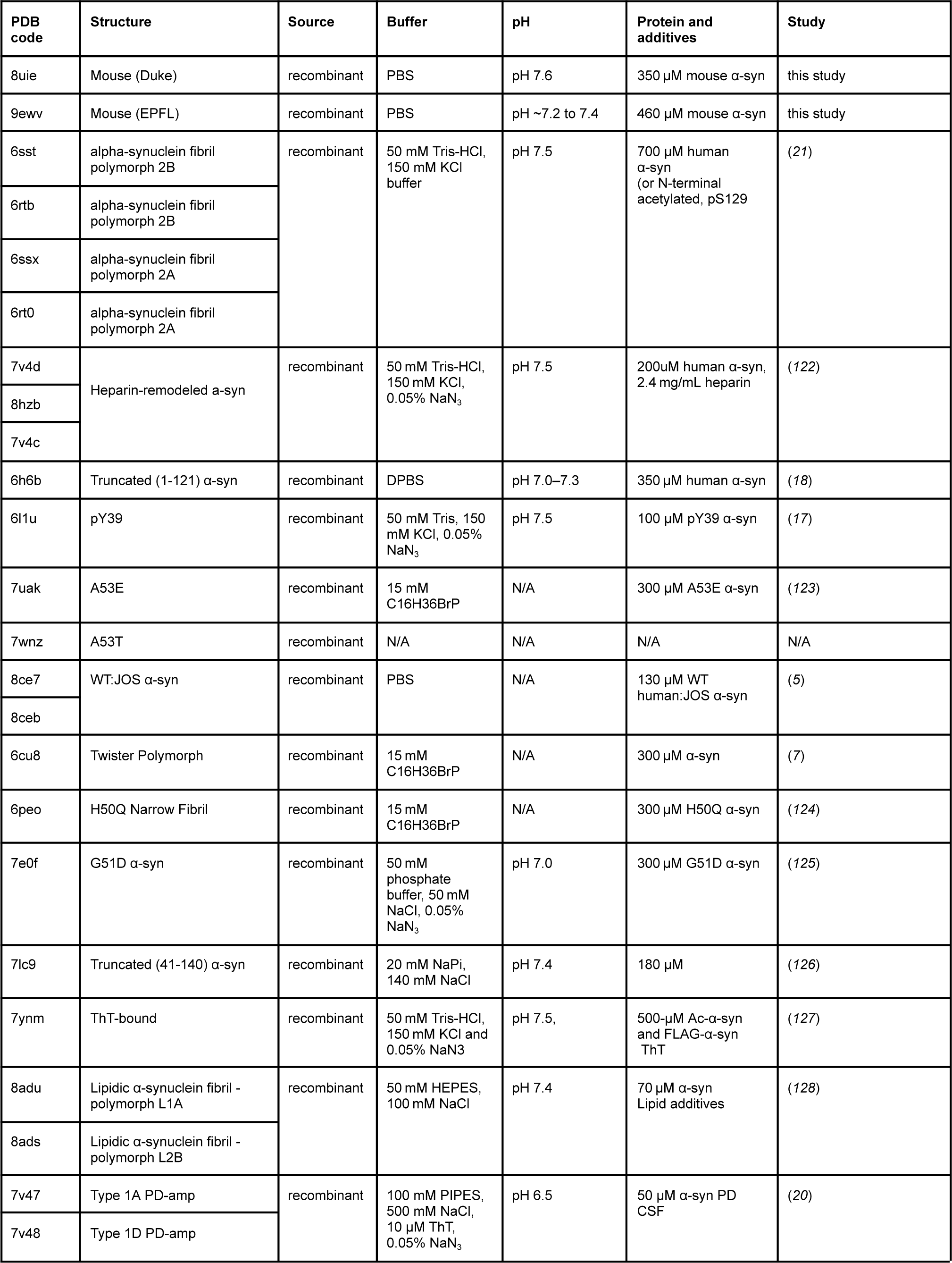
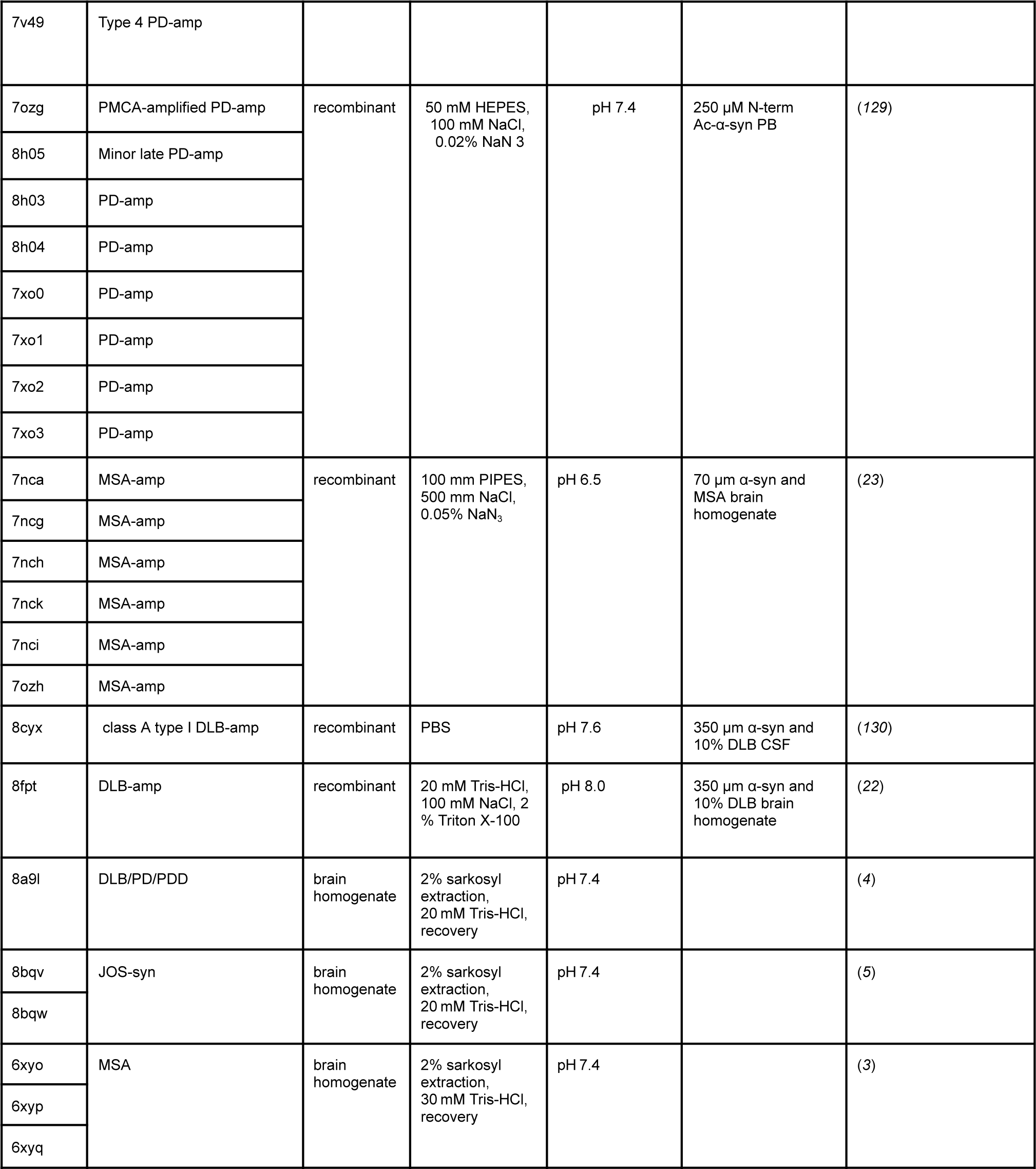
Fibril generation conditions.

**Fig. S1.**
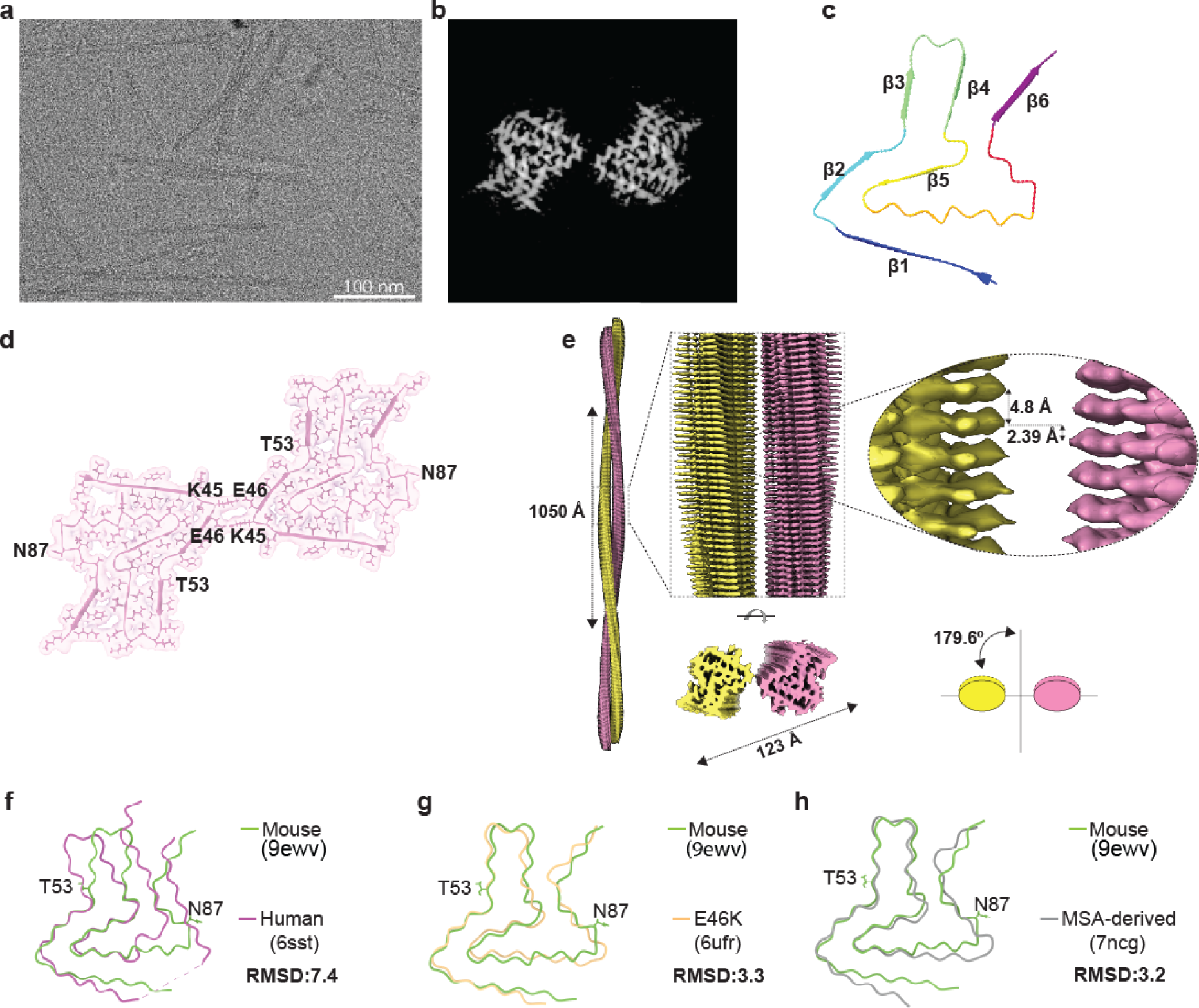
Cryo-EM structure and atomic models of EPFL mouse α-syn fibrils. (a) Representative cryo-EM micrograph of mouse α-syn fibrils with an indicated scale bars of 100 nm. (b) Cross-sectional view of the cryo-EM map density with detailed internal structure of a mouse α-syn fibril. (c) Cartoon representation of the fitted atomic model with indicated six β-strands (β1-β6), each delineated by a distinct color. (d) Cryo-EM density map with fitted atomic model of mouse α-syn spans residues 34-97, highlighting key residues K45 and E46 implicated in the formation of a salt bridge between protofilaments and contributing to the stabilization of the fibril structure. Additionally, residues N87 and T53 are highlighted, noting their variation from the human α-syn sequence. (e) Representation of overall mouse α-syn fibril architecture, noting a mean crossover distance of 1050 Å with a width of 123 Å, exhibiting pseudo-2fold symmetry with a rise per subunit of 2.39 Å and a twist of 179.6° highlighted. Global fit comparative analysis of the mouse α-syn fibril (9ewv, green) structure and (f) recombinant human α-syn fibril atomic model (6sst, magenta), (g) recombinant E46K (6ufr, yellow), and (h) MSA-amplified (7ncg, gray) with visualized T53 and N87 amino acid schematics and indicated aligned total RMSD values.

**Fig. S2.**
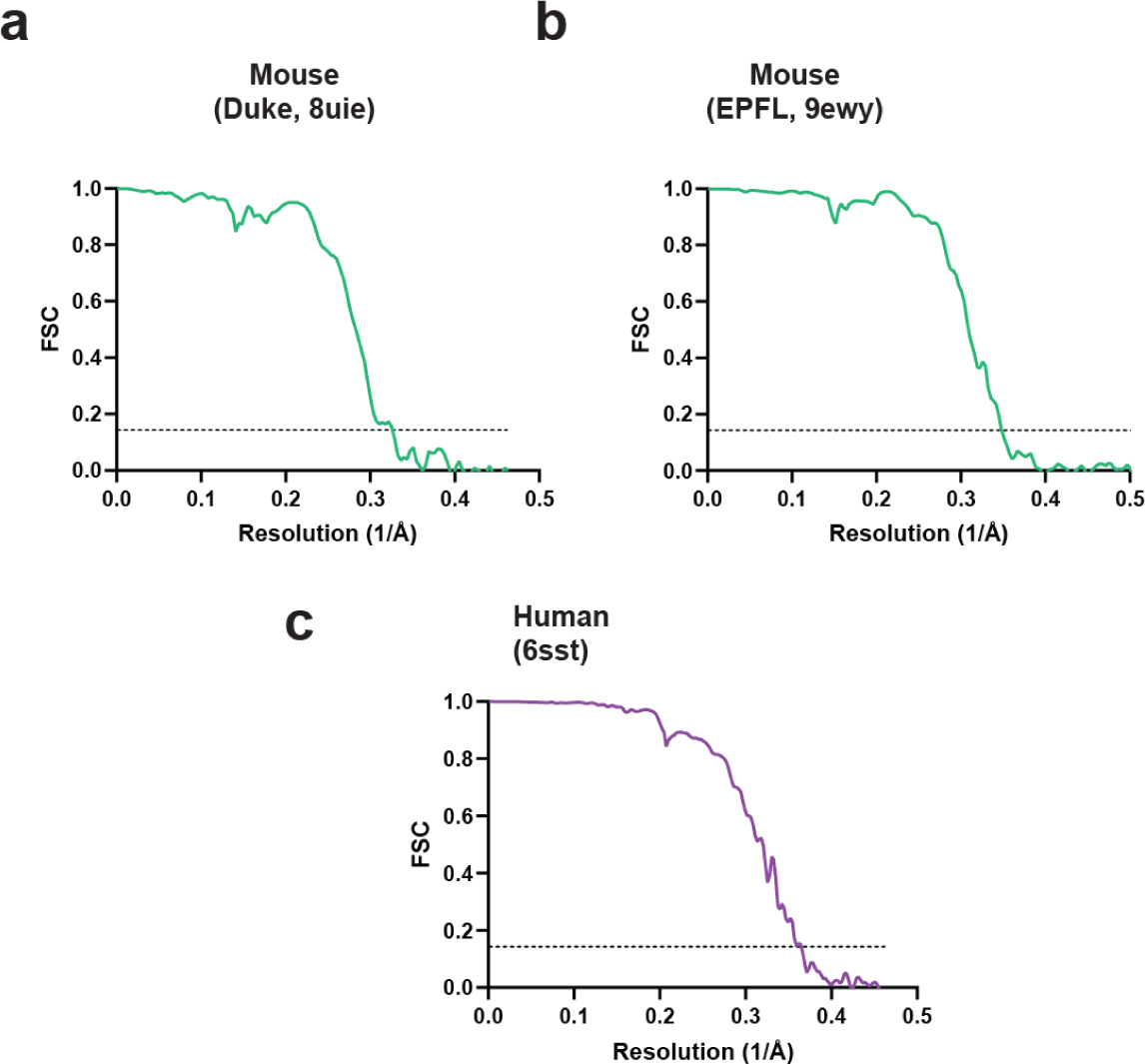
High resolution estimation of the cryo-EM map of mouse and human recombinant α-syn fibrils. Fourier shell correlation (FSC) resolution estimation and validation for the 3D reconstruction of the cryo-EM collected images of procured mouse fibrils generated by Duke (a) and EPFL (b) research groups. (c) FSC estimation plot of human α-syn fibrils collected and generated at Duke University.

**Fig. S3.**
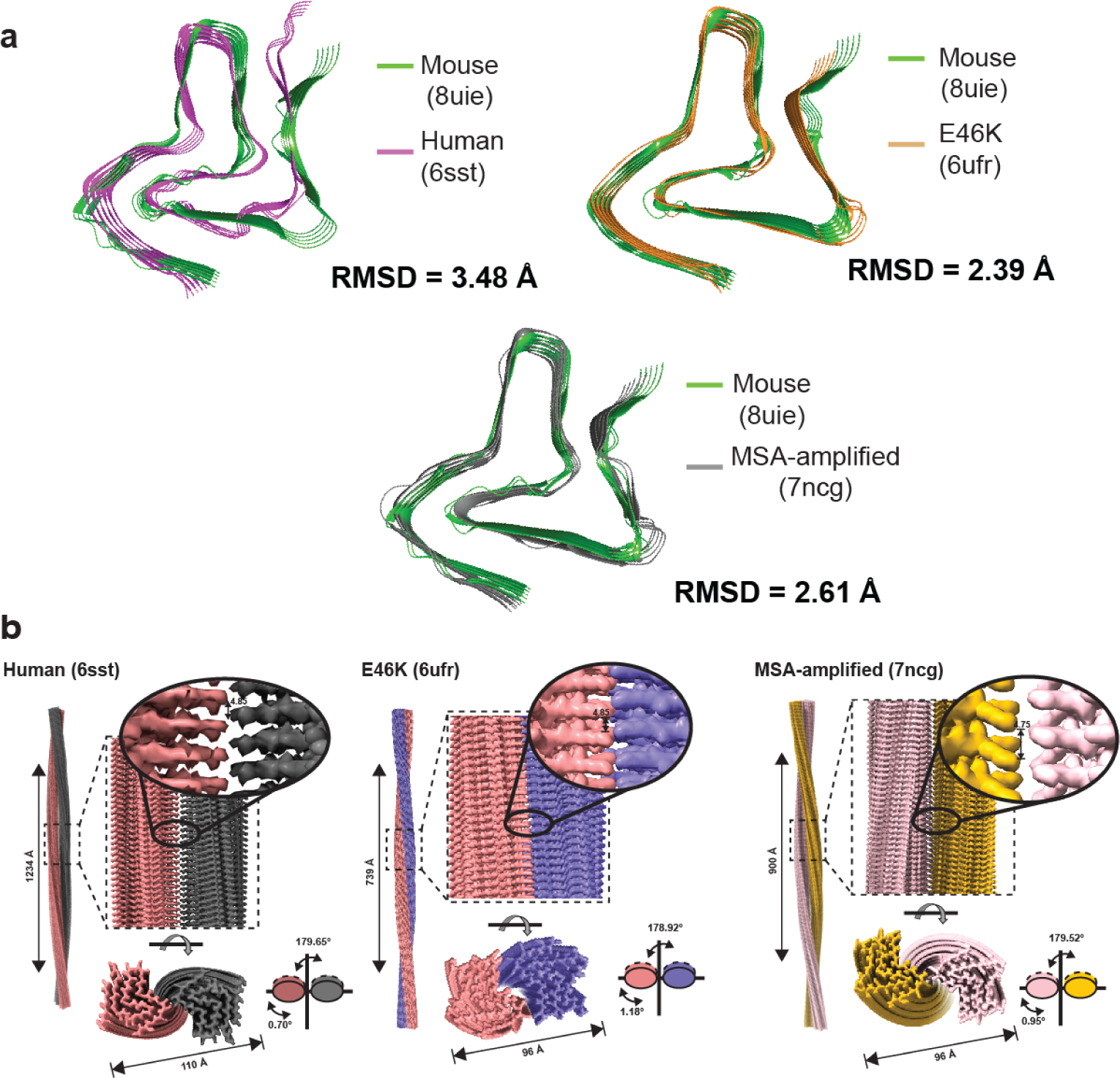
The distinct arrangement of mouse β-sheets aligns with E46K and MSA-amplified structures. (a) Overlay of human, E46K, and MSA-amplified structures compared with the mouse α-syn fibril structure generated at Duke (8uie), shown as individual RMSD values for each amino acid. The alignment was based on a global fit comparison. (b) Cryo-EM 3D reconstruction density map of the human (6sst), E46K (6ufr), and MSA-amplified (7ncg) α-syn fibrils, with a cross-section view. The pitch length, helical rise, and twist angle are highlighted.

**Fig. S4.**
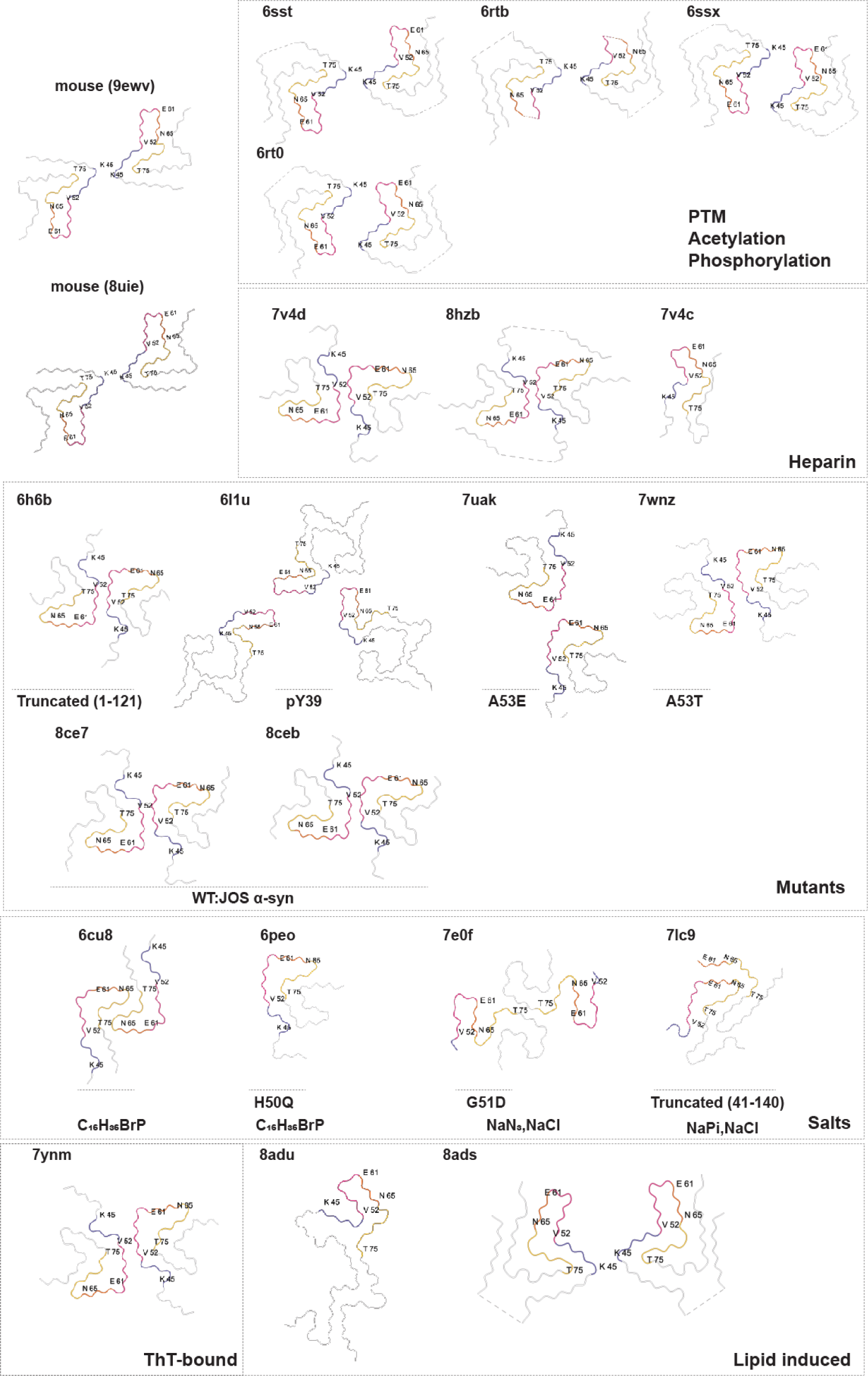
Summary of previously resolved recombinant α-syn fibril structures in relation to mouse α-syn fibril Cryo-EM maps. Cryo-EM maps of mouse α-syn fibrils generated from EPFL (9ewv) and Duke (8uie) research groups with β-sheets color-coded: β2 (blue), β3 (red), β4 (orange), and β5 (yellow). Previously recorded structures are categorized based on the unique characteristics that were employed for fibril generation. These characteristics are elaborated in Table S2. The color scheme is consistently maintained across the entire structure panel, serving to emphasize the structural features that are specific to the mouse α-syn structures.

**Fig. S5.**
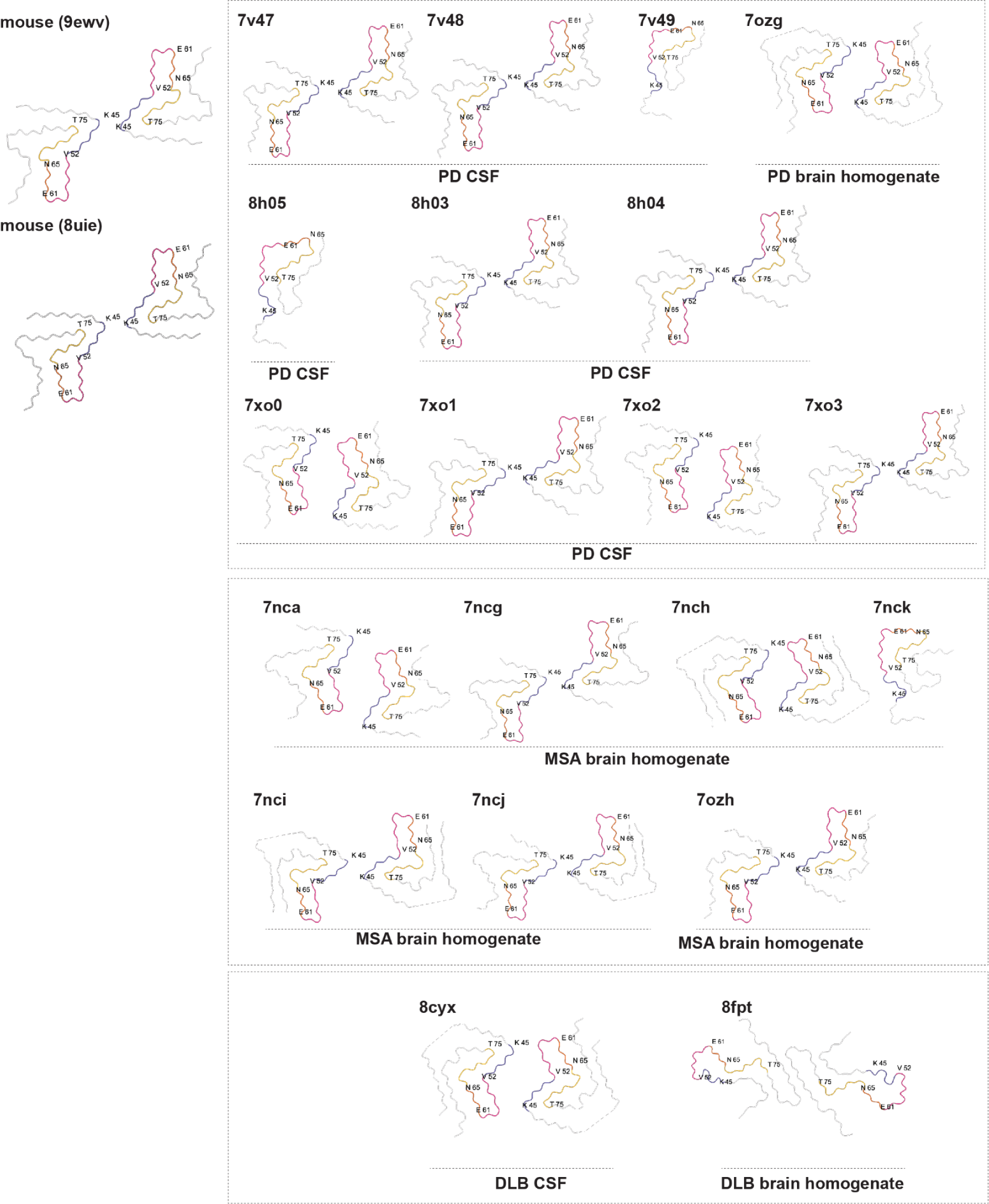
Structures of recombinant α-syn fibrils amplified from patient material in comparison to mouse α-syn fibril Cryo-EM maps. Cryo-EM maps of mouse α-syn fibrils curated at Duke (8uie) and EPFL (9ewv) with β-sheets color-coded as β2 (blue), β3 (red), β4 (orange), and β5 (yellow). Previously reported structures include α-syn fibrils generated in presence of patient material including brain homogenates and cerebrospinal fluid and detailed in Table S2.

**Fig. S6.**
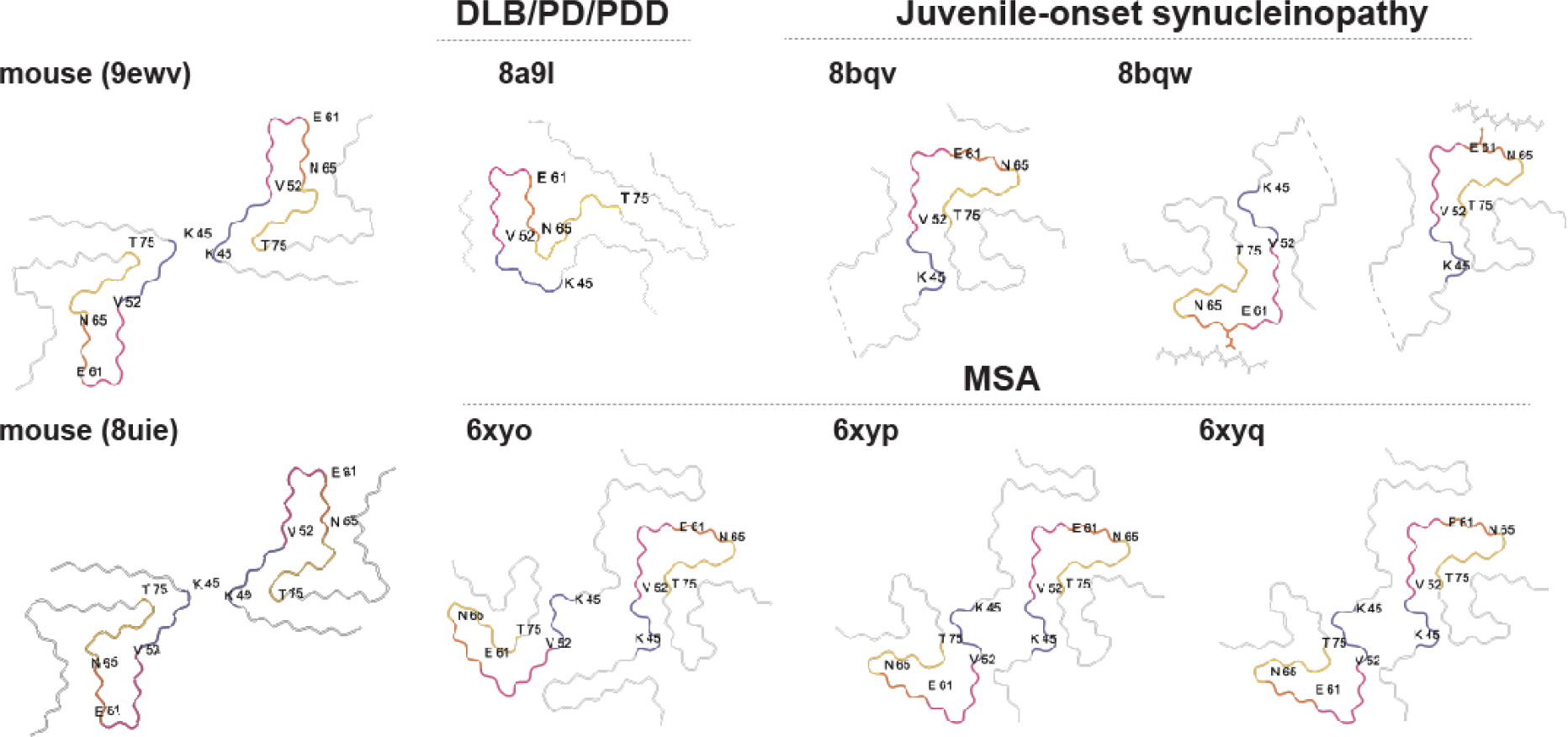
Structures of *ex vivo* α-syn fibrils extracted from brain homogenates in comparison to mouse α-syn fibril Cryo-EM maps. Cryo-EM maps of mouse α-syn fibrils curated at Duke (8uie) and EPFL (9ewv) with β-sheets color-coded as β2 (blue), β3 (red), β4 (orange), and β5 (yellow). Previously reported structures include ex vivo α-syn fibrillar assemblies extracted from patient material and detailed in Table S2.

**Fig. S7.**
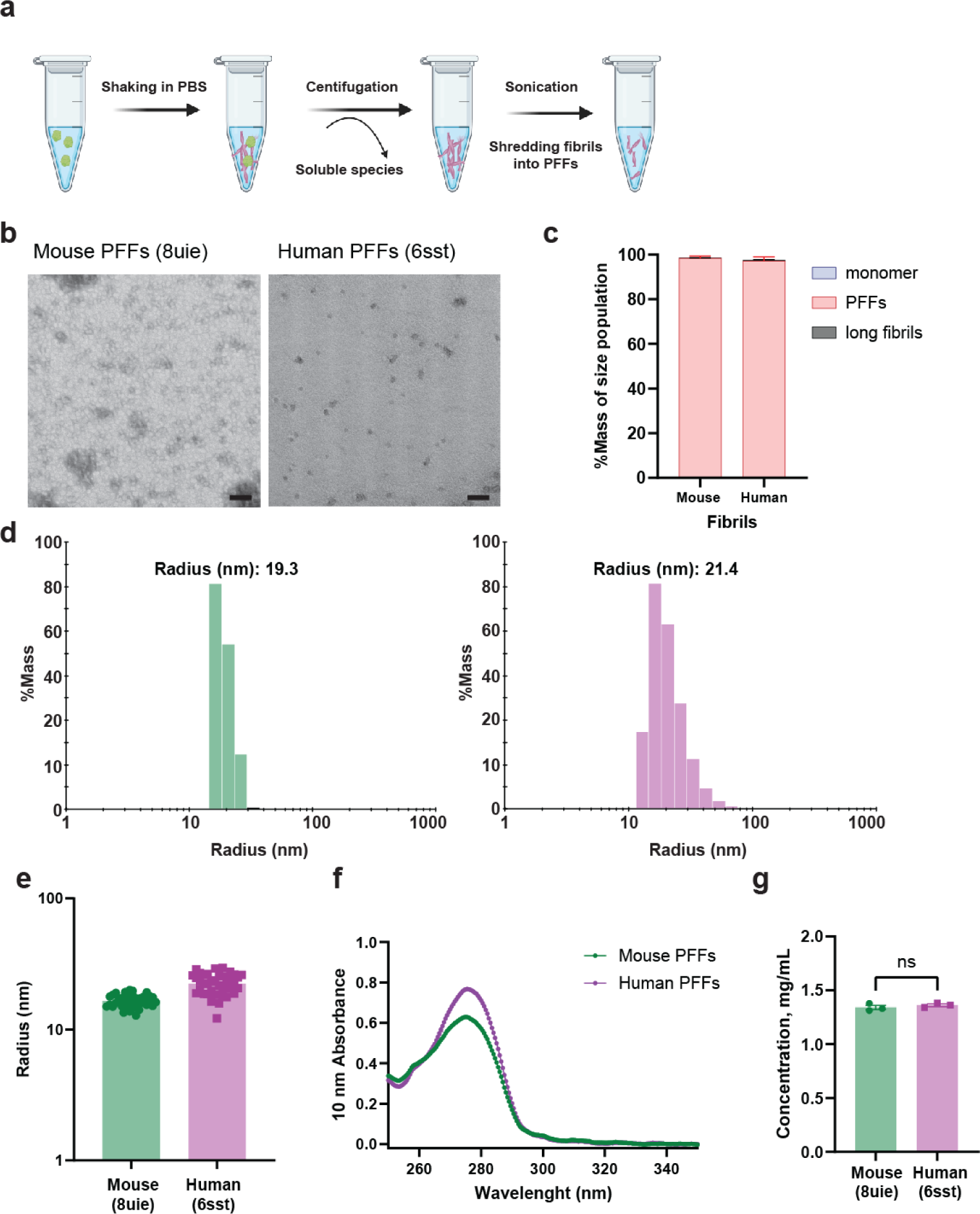
Generation and validation of mouse and human α-syn PFFs. (a) Schematic representation of the α-syn PFF preparation process, which includes the initial formation of aggregates through continuous shaking in PBS, separation of large aggregates using multiple cycles of high-speed centrifugation, and sonication to achieve a uniform size of short fibrils or PFFs. (b) Transmission electron micrographs of mouse α-syn (left) and human (right) PFFs after extraction and sonication cycles. (c) Group analysis of the size population proportions of sonicated mouse and human α-syn preparations, (d) representative radius distribution relative to percent of mass of mouse α-syn, (e) representative average radii of human α-syn,(f) UV absorbance spectra and (g) coefficient extinction adjusted concentration. Each data point or S.E.M in panel c, e and g are extracted from a single acquisition from three independent experiments with ten measurements analyzed per group. Each dot in panel f is the mean of two technical replicates from three independent batches. Significance was assessed via 2-tailed t-tests with ns for not significant.

**Fig. S8.**
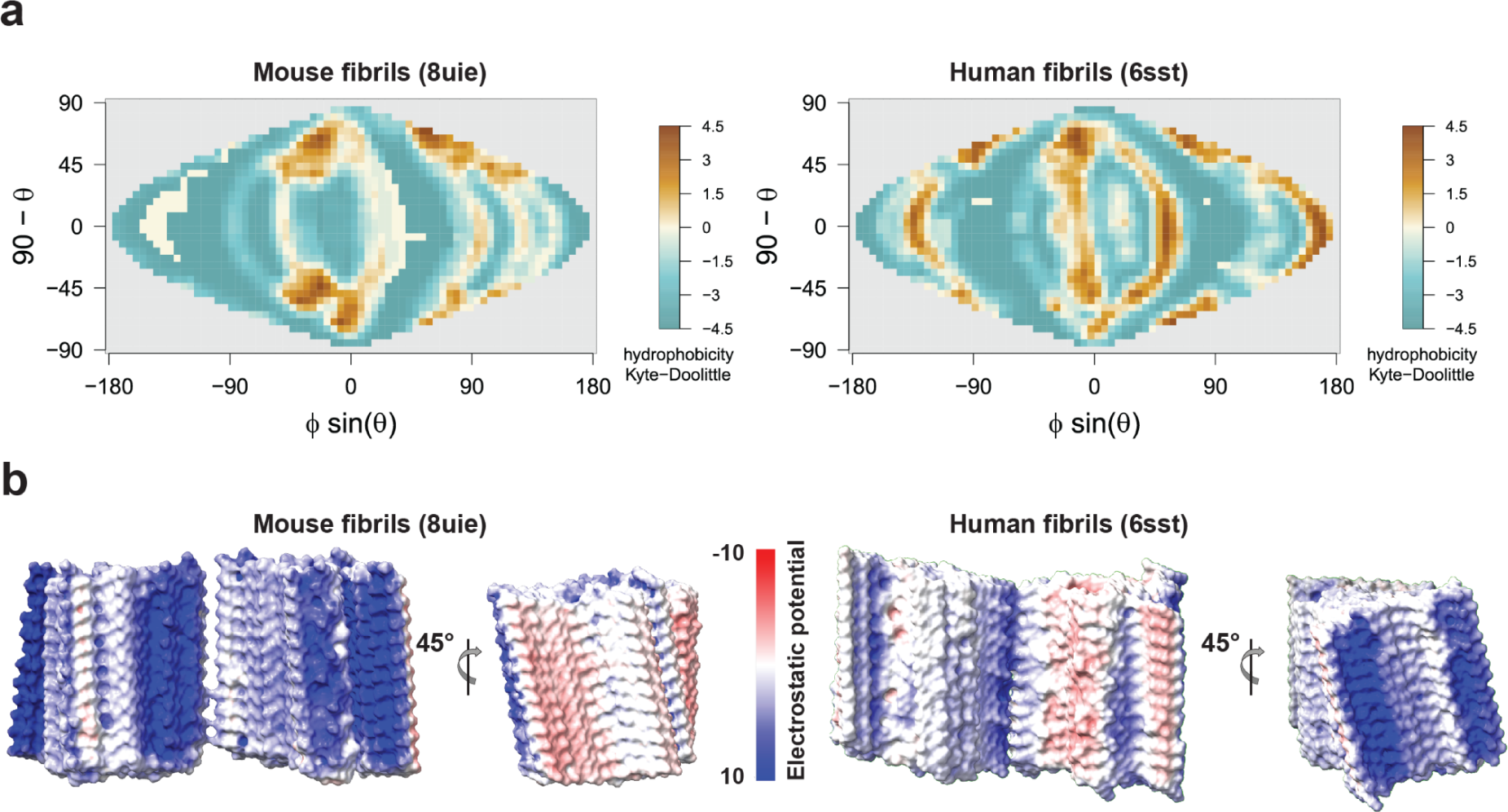
Mouse α-syn fibrils exhibit distinct characteristics of hydrophobicity and electrostatic surface charge. (a) Molecular 2D KD hydrophobicity map projecting 24 α-syn fibril layers of β-stacks with spherical coordinates (φ, θ) and the Sanson-Flamsteed 2D projection. The projected 2D map is divided into a grid of 36 × 72 cells. Each cell is smoothed by averaging its value with those of the eight surrounding cells, and is associated with the average of the corresponding hydrophobicity values. (b) Cross and side views of the electrostatic potential maps of mouse and human α-syn fibrils, generated using molecular dynamic simulations of atomic models placed under physiological pH and salt conditions.

**Fig. S9.**
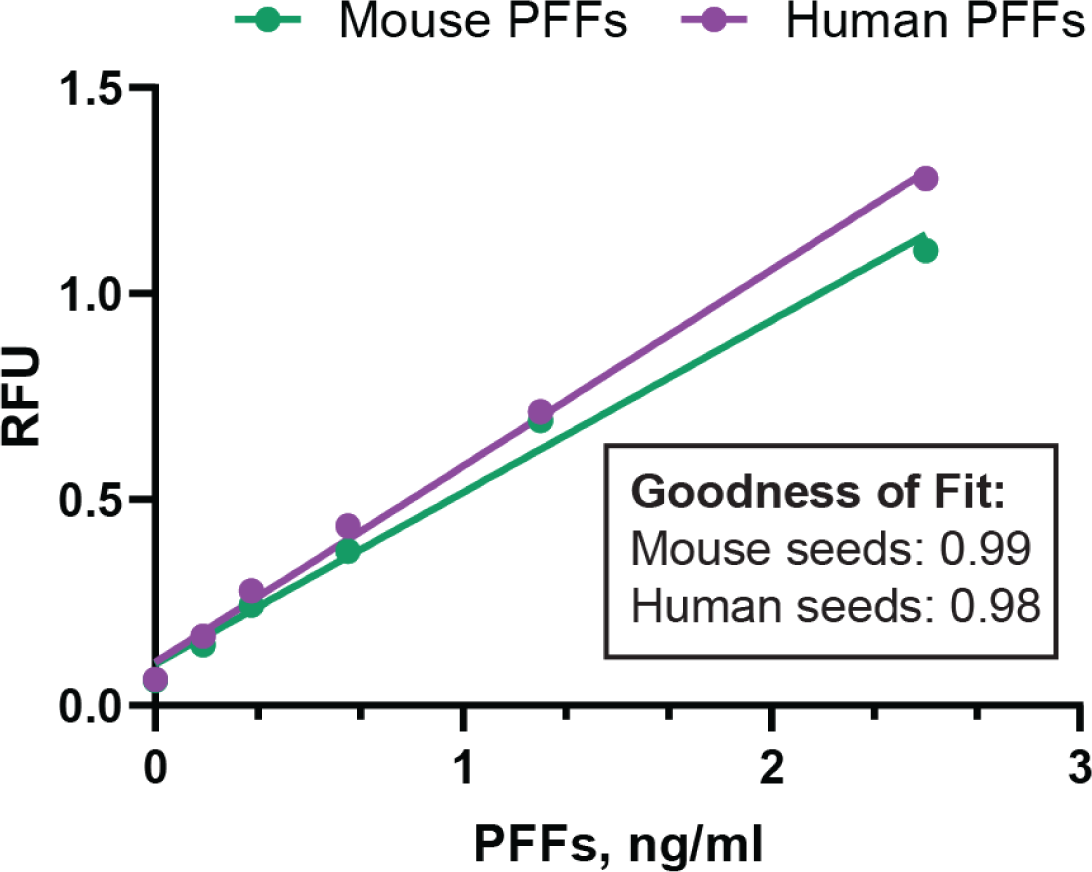
Evaluation of α-syn ELISA using control recombinant mouse and human α-syn fibril PFFs. Standard curve generated from mouse and human PFFs in a pan-α-syn aggregate ELISA. The ELISA was utilized to quantify the level of aggregates present in the cell lysates. The standard curve, along with the indicated goodness of fit and corresponding r² values, provides a reliable measure for interpreting the results. Each data point represents the mean from three technical replicates from two independent experiments with errors bars indicating S.E.M.

**Fig. S10.**
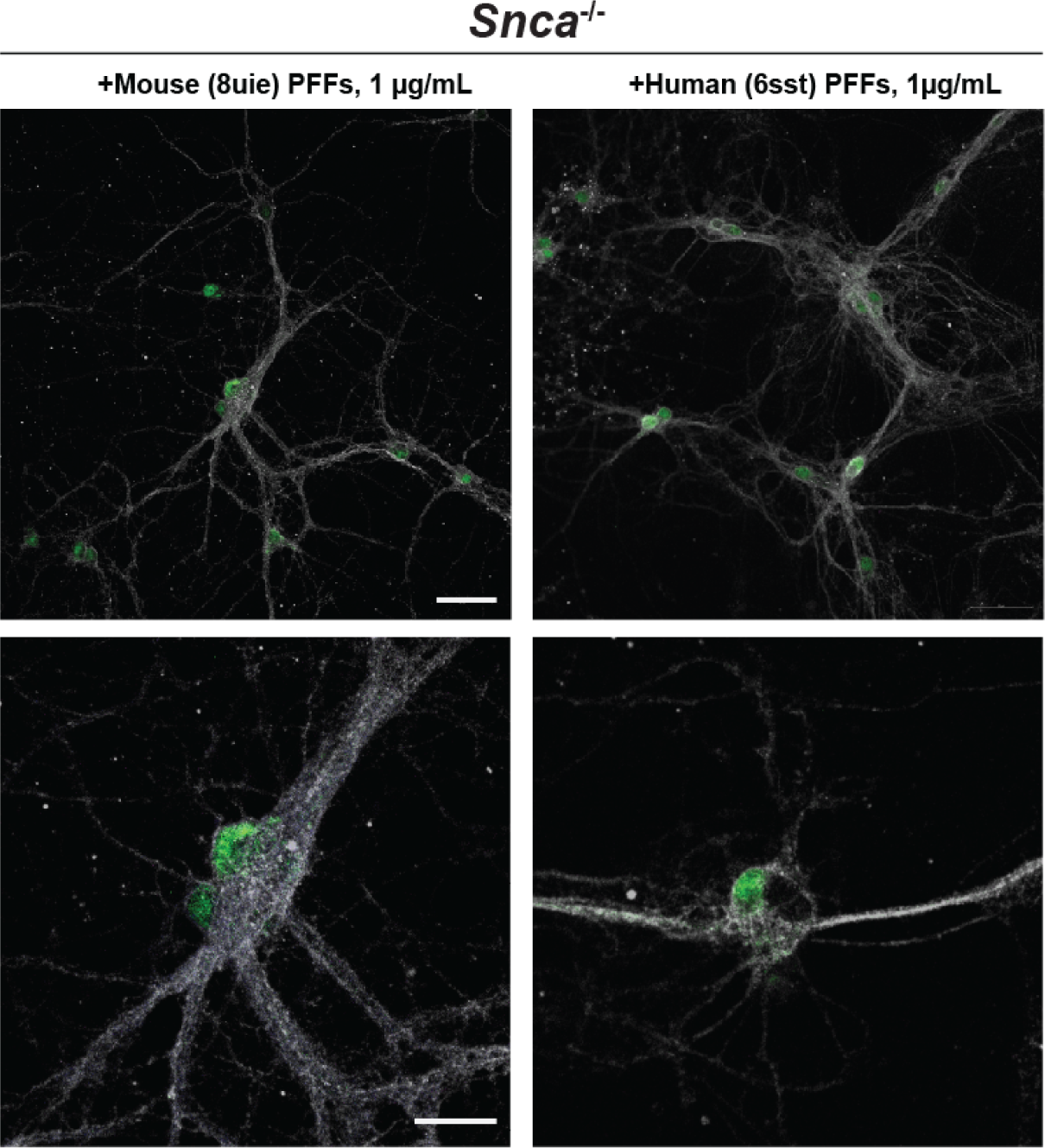
Seeding propensity following α-syn PFF treatment in *Snca*^−/−^ hippocampal neuronal culture. Representative Immunostaining of mouse *Snca*^−/−^ primary hippocampal neurons treated with mouse or human α-syn PFFs for 14 days before fixing. Magenta indicates pS129-α-syn staining, grey color indicates tau and green as NeuN. The scale bar represents 50 µm for the main images and 20 µm for the magnified images.

**Fig. S11.**
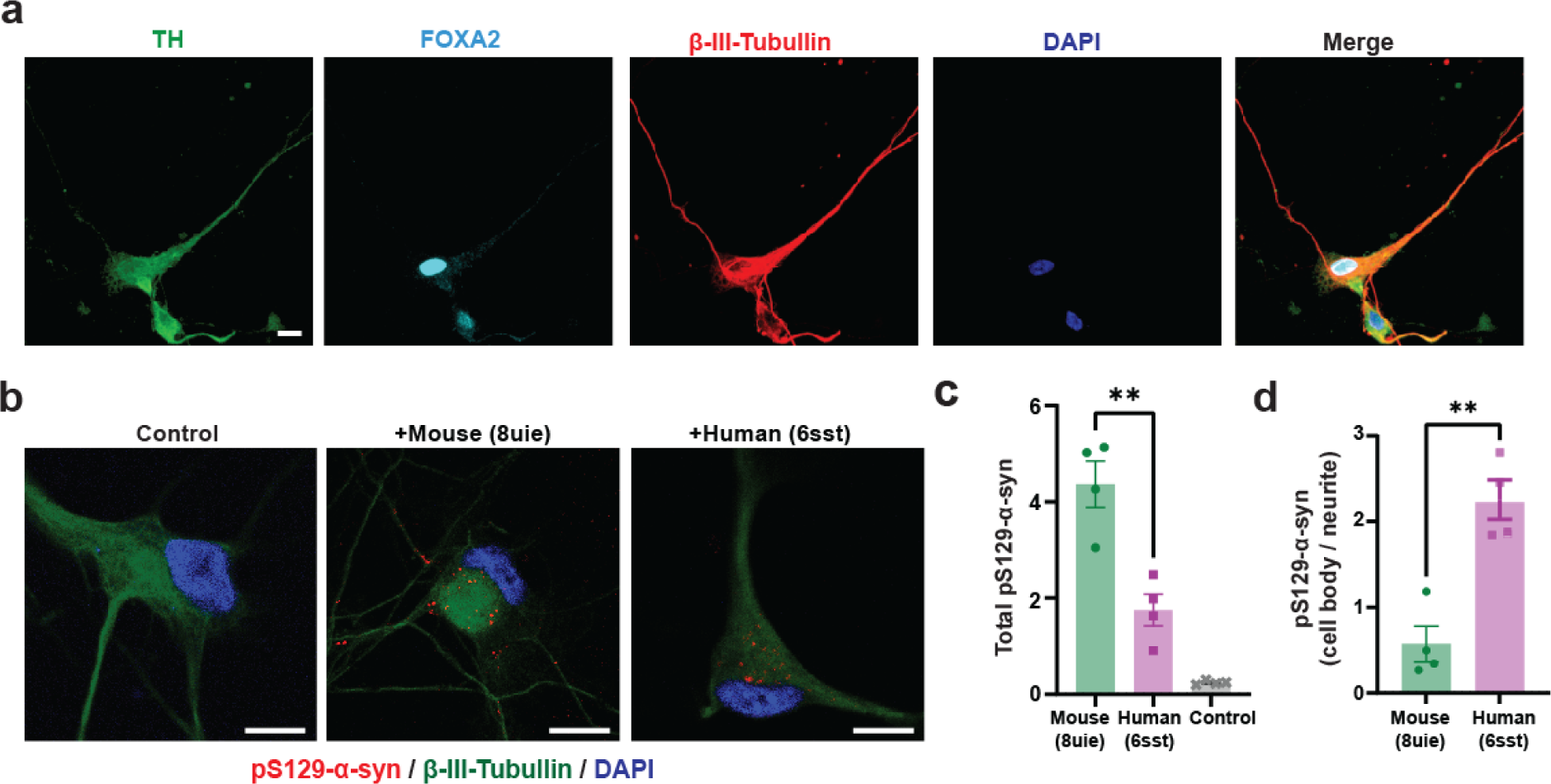
Elevated p-S129-α-syn in iPSC-derived dopaminergic neurons followed the treatment with mouse α-syn PFFs. (a) Representative Immunofluorescence images of ∼70-day-old iPSCs-derived DA neurons stained with the neuronal marker β-III-tubulin (red), dopaminergic markers tyrosine hydroxylase (TH, green) and FOXA2 (cyan) counterstained with DAPI (blue). The scale bar is 10 μm. (b) Representative immunostaining of iPSCs-derived DA neurons treated with either Alexa-568 labeled mouse or human α-syn PFFs at 10 μg per mL for 7 days, along with intact control cells. iPSCs-derived DA neurons were fixed 7-days post initial treatment, permeabilized, and stained for β-III-tubulin (green) to outline neuronal architecture, pS129-α-syn (red), and DAPI (blue). The scale bar is 5 μm. (b) Levels of pS129-α-syn in the β-III-tubulin area in iPSC-derived DA neurons after 7 days of treatment with 10 μg/mL of PFFs or control. (c) The quantity of pS129 puncta relative to the β-III-tubulin area measured in each image and compared across conditions. (d) Proportion of abundance of pS129-α-syn puncta localized in cell bodies and neurites in group analysis between mouse and human α-syn PFF treatments. Each data point in c and d represent the mean value of the images from one well (n=4) and errors bars indicate S.E.M with **p<0.01 from 2-tailed t-tests.

## Notes

### Competing Interest Statement

The authors have declared no competing interest.

